# Discrete Morse Graph Construction for High-Dimensional Transcriptomic Data

**DOI:** 10.64898/2026.09.22.753642

**Authors:** Lucas Magee, Rohan Gala, Kyle Travaglini, Uygar Sümbül, David R. Haynor, Samantha Chen, Tristan Brugère, Yusu Wang, Partha P. Mitra, Michael Hawrylycz

## Abstract

Single-cell transcriptomic atlases resolve thousands of putative brain cell types, yet common analytical workflows often rely on low-dimensional embeddings that can distort neighborhood relationships and obscure continuous variation between closely related populations. We introduce single-cell Discrete Morse Graph Construction (scDMGC), a deterministic framework based on topological data analysis and discrete Morse theory that extracts a compact graph from neighborhood structure in high-dimensional gene-expression space. Because graph edges are derived from the original *k*-nearest-neighbor complex, scDMGC provides an interpretable multiscale representation of transcriptomic organization without requiring a low-dimensional embedding. Local maxima represent coherent transcriptional states, saddles quantify their separation, and gradient paths capture intermediate states and continuous transitions. Applied to cortical and hippocampal datasets, scDMGC recovers established inhibitory-neuron classes and elucidates transcriptional gradients. In mouse whole-cortex data, Morse graph structure evaluates cell-type hierarchy and type robustness. Persistence across scales provides a quantitative measure of cell-type identity and discrete versus continuous relationships. Finally, in Alzheimer’s disease single-nucleus data, scDMGC distinguishes changes in cell-type abundance from disease-associated shifts in transcriptional state. scDMGC therefore provides a multiscale framework for quantifying whether transcriptional populations form persistent cell types, how strongly external annotations are supported by intrinsic data structure, how cell types relate to one another, and where discrete organization transitions into continuous variation.

## Introduction

Single-cell and spatial transcriptomics have generated large, high-dimensional datasets that are transforming the construction of brain cell atlases across species.^1–4^ Single-cell and single-nucleus RNA sequencing (sc/snRNA-seq) are now routinely used to characterize anatomical and transcriptomic organization,^1–5^ developmental programs,^6^ evolutionary variation,^6,7^ disease-associated changes,^8^ and other biological processes. A central computational challenge is to extract reproducible structure from these data and relate transcriptional variation to anatomy, cell identity, and biological condition.^9^

Most analyses address the high dimensionality of transcriptomic data through feature selection and dimensionality reduction.^10–12^ Methods such as principal component analysis, t-distributed stochastic neighbor embedding (t-SNE)^13^, and uniform manifold approximation and projection (UMAP) ^14^ produce compact representations that facilitate visualization, clustering, and interpretation of cellular heterogeneity. These approaches have been commonly used in developing and interpreting modern brain cell-type taxonomies.^1–5^ However, strong dimensionality reduction necessarily alters aspects of the geometry of the original data. Distances and neighborhood relationships can be substantially distorted,^13,15^ and neither local nor global metric structure is guaranteed to be preserved.

This limitation becomes particularly important when the biological structure of interest extends beyond well-separated clusters. Transcriptomic variation can contain both discrete organization associated with relatively stable cell identities and continuous structure associated with anatomical gradients, developmental trajectories, physiological states, and transitions between closely related types.^16,17^ Distinguishing these forms of variation requires methods that can identify robust local concentrations of cells while simultaneously representing the relationships and intermediate states connecting them. Low-dimensional embeddings can reveal such structure qualitatively, but quantitative conclusions about boundaries, gradients, or similarity between nearby populations may depend on the projection itself.^13,15^ This motivates methods that operate directly on neighborhood relationships in the selected high-dimensional gene-expression space while producing a compact and interpretable representation.

Topological data analysis (TDA) provides a mathematical framework for this problem.^18–20^ Rather than requiring a particular low-dimensional coordinate system, TDA characterizes structural features of data through connectivity and topology across scales. For discrete datasets, local relationships can be represented by simplicial complexes composed of vertices, edges, triangles, and higher-dimensional simplices. Persistence provides a principled mechanism for distinguishing prominent structure from features that arise only over a narrow range of scales, thereby reducing sensitivity to sampling variability and noise.

Morse theory provides a complementary description of structure in terms of critical points and the gradient paths connecting them.^21,22^ In a continuous landscape, maxima, minima, and saddles organize the gradient flow and define the relationships among prominent features. *Discrete Morse theory* (DMT), with the seminal work of Forman,^22^ extends these concepts to combinatorial complexes, allowing analogous structures to be extracted from discrete data. DMT and related approaches have been applied to problems including image skeletonization^21,23^ and neuronal morphology reconstruction,^24–26^ but their application to large, high-dimensional single-cell transcriptomic datasets has remained limited.^20^

Here we introduce single-cell *Discrete Morse Graph Construction* (scDMGC), a framework that combines discrete gradient structure with persistence-based simplification to extract a compact graph directly from high-dimensional scRNA-seq data. Building on recent theoretical and computational advances in discrete Morse graph reconstruction,^27^ scDMGC constructs a one-dimensional graph skeleton from a local neighborhood complex while simultaneously defining a gradient field that associates each cell with a representative on the graph. Local maxima identify candidate transcriptional types, saddles characterize the separation between neighboring maxima, and gradient paths connect them along transcriptional ridges containing cells with intermediate profiles. Persistence provides a natural scale parameter for determining which peaks and connections remain prominent as progressively weaker structure is removed.

Importantly, scDMGC does not require an intermediate two- or three-dimensional embedding to define this structure. Although the resulting Morse graph provides a compact representation that can be readily visualized, its paths are constructed from neighborhood relationships in the selected gene-expression space. The framework therefore provides complementary information to conventional clustering and visualization methods: clusters can be represented as persistent peaks, while saddles and connecting paths quantify the strength and structure of relationships between them. In this way, discrete cell identities and continuous transcriptional variation can be examined within a common representation.

We evaluate scDMGC across four biological settings. First, using mouse cortical transcriptomic data,^1,5^ we examine how Morse structure captures cell-type identity and the separation among established transcriptomic populations. Second, using GABAergic interneurons from hippocampal CA1,^28^ we reconstruct a previously described transcriptional continuum at single-cell resolution and characterize its internal organization. Third, we use persistence across scales to derive a graph-based organization of mouse GABAergic neurons and quantify the coherence and distinctness of externally assigned cell-type labels. Finally, using single-nucleus data from the Seattle Alzheimer’s Disease Brain Cell Atlas (SEA-AD),^8,29^ we examine disease-associated changes in the Morse landscape and distinguish changes in cell abundance from shifts in transcriptional state. Together, these analyses demonstrate how discrete Morse graph reconstruction can provide a multiscale representation of both discrete cell identity and continuous transcriptomic organization directly from high-dimensional single-cell data.

### 1. Discrete Morse graph construction for omics data

The scDMGC algorithm builds on topological data analysis and discrete Morse theory (DMT) (**Methods**).^18,19,21,22,30^ DMT operates on a simplicial complex endowed with a scalar function and provides a combinatorial analogue of critical points and gradient flows. Recent methods use this framework to reconstruct graph skeletons from sampled scalar fields by extracting unstable one-manifolds that follow prominent ridges. ^25–27^ Persistence-based simplification removes low-prominence features while retaining the dominant peaks, saddles, and connecting ridges. We adapt a recent discrete Morse graph-reconstruction algorithm^27^ for high-dimensional single-cell transcriptomic data.

Direct application of graph-reconstruction methods developed for lower-dimensional point clouds is computationally impractical at the scale of single-cell datasets. We therefore represent the data using a two-dimensional *k*-nearest-neighbor simplicial complex together with a Jaccard-based lower-star filtration (**Methods**), a data structure for capturing multiresolution. The complex contains cells as vertices, *k*-nearest-neighbor relationships as edges, and triangles induced by these relationships. Its simplices are ordered according to local neighborhood overlap. From this filtered complex, scDMGC computes a discrete gradient vector field and extracts a persistence-simplified Morse graph. Each cell follows the discrete gradient field to a unique vertex on this graph, which we refer to as its *Morse representative*.

**Fig. 1** illustrates the workflow using 57,902 somatostatin (SST) expressing cells from mouse cortex.^5^ Starting from a log-normalized cell-by-gene expression matrix, we select 2,500 highly differentially expressed genes to define the input expression space (**Fig. 1A**). A two-dimensional *k*-nearest-neighbor complex is then constructed using Euclidean distance, with cells represented as vertices, neighboring cell pairs as edges, and adjacent triples as triangles (**Fig. 1B**). For each edge (*u*, *v*), compute the Jaccard similarity *J*(*u*, *v*), defined as the fraction of neighbors shared by *u* and *v*. Each vertex is assigned the maximum Jaccard similarity *J*(*u*, *v*), over its incident edges (**Methods**). Thus, two cells have a high similarity when their local neighborhoods strongly overlap (**Fig. 1C**), and a cell with a high Jaccard index has many neighbors that are like it; such a cell is intuitively a good candidate for the archetype of a cell type.

**Figure 1.**
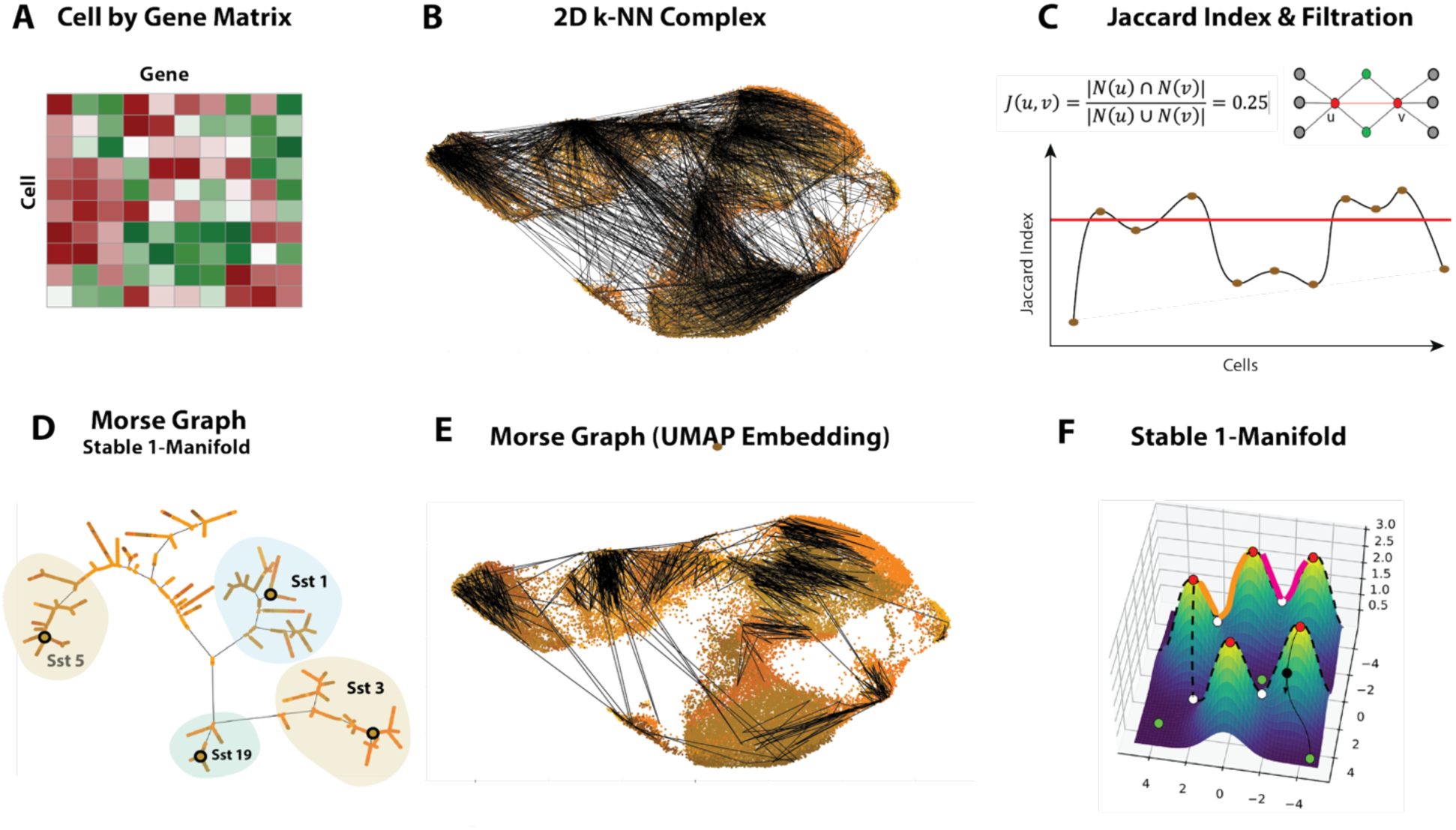
Discrete Morse graph construction. (A) Log-normalized cell-by-gene expression matrix. (B) Two-dimensional 25-nearest-neighbor simplicial complex built from *Sst* expressing cells in mouse cortex anterior cingulate area5 containing simplices of dimension at most 2: cells (vertices), edges, and triangles. Only edges shown in visualization. (C) Each edge is assigned a Jaccard similarity equal to the fraction of shared neighbors. The resulting vertex function orders the simplices to define the filtration used by scDMGC. (D) scDMGC extracts a Morse graph in the input gene-expression space, comprising peaks, saddles, and optimal gradient paths (unstable one-manifolds). Points shown are Morse maxima with annotated cell type labels from original clustering.5 (E) Projection of the *Sst* Morse graph onto a UMAP embedding illustrates the distortion of short-range relationships in two dimensions. (F) Conceptual landscape in which transcriptional states form peaks connected through saddles and gradient paths. Two peak–saddle–peak paths are highlighted in orange and red.

The Jaccard field defines the filtration used by scDMGC. Local maxima identify densely and coherently sampled transcriptional states, while saddles characterize the connections between neighboring maxima. Persistence quantifies the prominence of these features and provides a continuous scale for simplifying the resulting structure.^18^ scDMGC extracts a Morse graph consisting of persistent peaks and saddles connected by optimal discrete gradient paths (**Fig. 1D**). Rather than projecting all cells into a low-dimensional coordinate system, the method identifies a compact subset of cells and edges that summarizes the organization of the original neighborhood complex.

This distinction is important because nonlinear dimensionality-reduction methods can substantially alter metric and neighborhood relationships.^31,32^ In *Cck*-expressing hippocampal CA1 interneurons,^28^ pairwise distances in a t-SNE embedding correlated moderately with longer-range Euclidean distances in the input expression space (ρ = 0.60), but only weakly for local 15-nearest-neighbor pairs (ρ = 0.10; **Fig. S1**). Similarly, projecting a Morse graph onto a UMAP embedding shows that cells adjacent in the original *k*-nearest-neighbor complex can appear widely separated in two dimensions (**Fig. 1E**). Thus, low-dimensional visualization can obscure precisely the local relationships used to define transcriptional continuity.

By construction, every edge in the Morse graph is inherited from the original *k*-nearest-neighbor complex. The Morse graph is also intrinsically multiscale. At low persistence thresholds, it retains a larger number of peaks, paths, and cells; increasing the threshold progressively removes weaker features and contracts the representation to the most prominent peaks and ridges. Across persistence thresholds, cells are substantially more likely to have a nearest neighbor represented on the Morse graph than expected for randomly selected cell subsets of equal size, and this enrichment is robust to the choice of *k* (**Fig. S2A**). The Morse graph therefore provides a compact, multiresolution representation of local transcriptomic organization that can be interpreted as a landscape of transcriptional peaks connected through saddles and gradient ridges (**Fig. 1F**).

We next asked whether this graph structure captures biological information beyond that contained in the selected Morse cells themselves. Although GABAergic interneurons share broad developmental origins^33^ and exhibit weaker regional specialization than glutamatergic neurons,^34^ cortical and hippocampal regions retain distinguishable GABAergic transcriptional signatures.^1,34,35^ We generated ten independent samples of 500 *Vip*-positive cells from each of 18 cortical and hippocampal regions, yielding 180 Morse graphs (**Fig. S3**). Regional separability based on Morse-cell subsets alone was like that obtained from random subsets of the same size (0.88 versus 0.85). When Morse graph connectivity was incorporated, separability increased to 0.95. Thus, the topology of the reconstructed graph contains regional transcriptional information that is not explained simply by which cells are selected. Graph comparisons were quantified using optimal-transport and Weisfeiler–Lehman-based distances (**Methods**).^36,37^

#### Discrete Morse graph construction and cell-type identity

High-throughput profiling and computational advances now permit systematic classification of neuronal transcriptomic diversity^1,4,17,28,34,38^ A transcriptomic cell type can be viewed as a set of cells with similar and distinguishable expression profiles.^17^ In scDMGC, such cells tend to share neighbors in the *k*-nearest-neighbor graph and therefore generate locally high Jaccard similarity (**Methods**). We define a *Morse cell type* as a vertex corresponding to a local maximum of the Jaccard-index field at a given persistence level. Distinct neighboring types produce an overall peak–saddle–peak structure, in which the reduction in Jaccard similarity between peaks quantifies their separation and the optimal gradient path identifies a direct transcriptional transition between them. Importantly, the Jaccard index need not decrease monotonically from a peak to the saddle or increase monotonically from the saddle to the neighboring peak; local fluctuations can occur along the discrete path, while persistence captures the overall prominence of the peaks relative to the intervening saddle. Edges with weak neighborhood overlap are excluded from the Morse graph.^27^ The prominence of each peak relative to its connecting saddle therefore provides a scale-dependent measure of evidence for cell-type distinctness.

To illustrate this interpretation, we analyzed two somatostatin-positive clusters, *Sst* 88 (2,775 cells) and *Sst* 91 (1,515 cells), identified in the mouse cortex.^5^ Both express the neuromedin B receptor gene *Nmbr* and arise from the medial ganglionic eminence. *Sst 91* is strongly associated with synaptic structure and organization (FDR q < 1.88 *x* 10^−8^), while *Sst88* more weakly associated with ion-channel activity (q < 8.93 *x* 10^−3^).^39^ **Fig. 2A** shows these clusters within a UMAP representation of 45,467 *Sst* cells. At the selected persistence threshold, the Morse graph *M* contained 239 vertices, including local maxima labeled *Sst* 88 and *Sst* 91 (**Fig. 2B**). We examined the optimal path *L* connecting these maxima (|*L*| = 44; **Fig. 2C**). Along this path, Jaccard index exhibited an overall decline from the *Sst* 91 peak toward an intervening saddle, followed by an increase toward the *Sst* 88 peak, with local fluctuations reflecting the discrete neighborhood structure of the data. This peak–saddle–peak profile defines a transition between the two transcriptional neighborhoods. Embedding the same path in the two-dimensional UMAP representation (**Fig. 2A**) illustrates how projection can distort local distance relationships that are retained in the original neighborhood graph.

**Figure 2.**
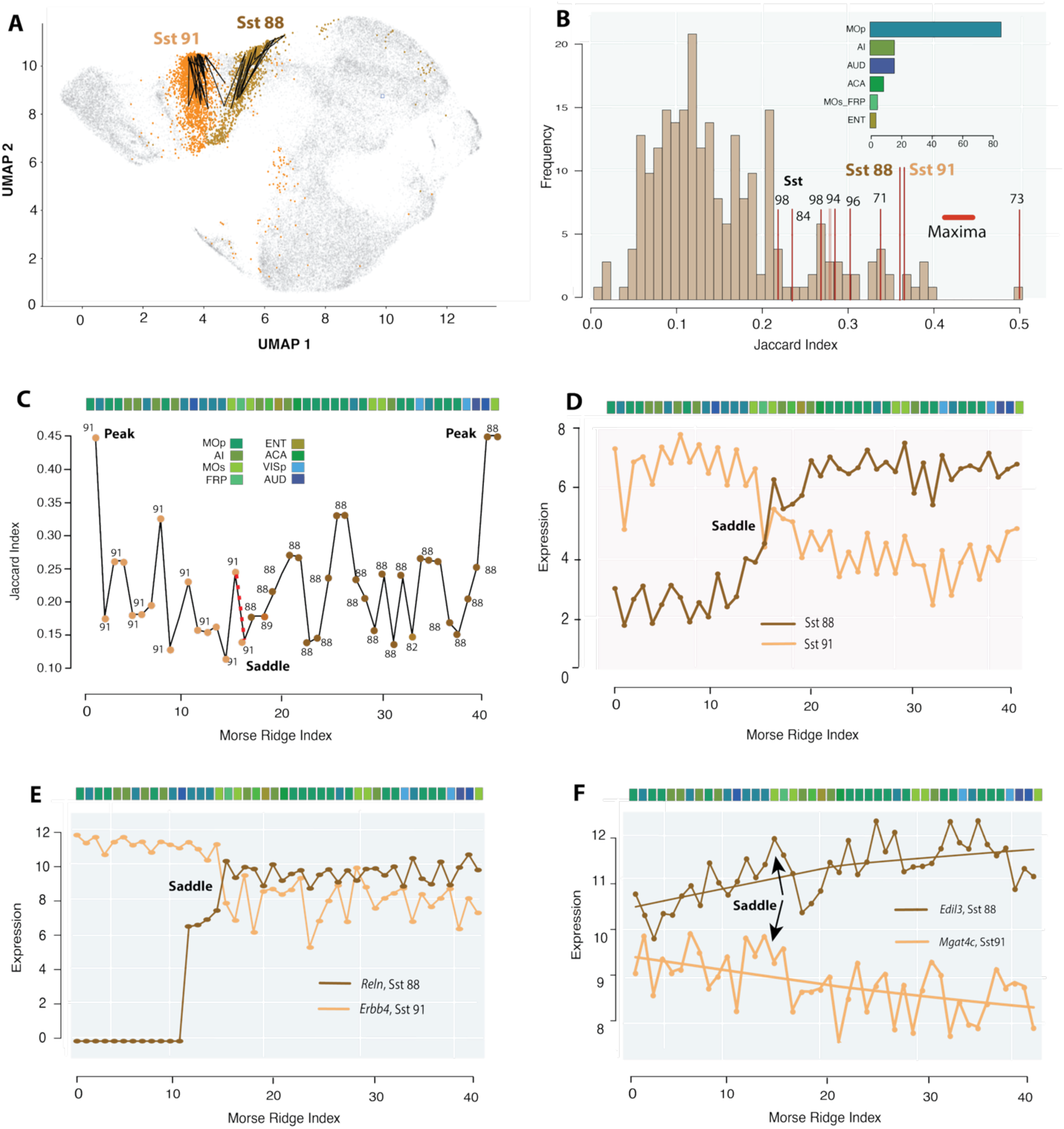
scDMGC resolves closely related cell types. (A) UMAP projection of *Sst* cells from anterior cingulate area,5 highlighting *Sst* 88 (orange), *Sst* 91 (brown), and a Morse path computed in the input gene-expression space and projected onto the UMAP. (B) Distribution of Jaccard-index values for the 239 vertices of the Morse graph; selected local maxima are labeled by *Sst* type. Inset shows the anatomical distribution of Morse vertices. (C) Morse path *L* (|*L*| = 44) connecting local maxima labeled *Sst* 91 and *Sst* 88. The dashed red segment marks the saddle edge (*L*_15_, *L*_16_) separating the two peaks. Jaccard similarity exhibits an overall peak–saddle–peak profile, with local fluctuations along the discrete path. (D) Mean expression trajectories of the 50 most differentially expressed genes between *Sst* 91 and *Sst* 88, showing a crossover near the saddle. (E) *Reln* and *Erbb4* show opposing expression transitions along the path. (F) *Edil3* and *Mgat4c* vary more gradually. Anatomical abbreviations: cortical regions Mop (primary motor area), AI (agranular insular area), AUD (auditory areas), ACA (anterior cingulate area), Mos (secondary motor area), FRP (frontal pole), and ENT (entorhinal area), VISp (primary visual).

The mean expression trajectories of the 50 most differentially expressed genes between *Sst* 88 and *Sst* 91 crossed near the saddle of *L* (**Fig. 2D**), consistent with this region marking the transition between the two expression profiles. The neuronal development and signaling gene *Reln*^40^ showed one of the strongest transitions, whereas *Erbb4*, a receptor tyrosine kinase implicated in several neurological disorders,^41,42^ changed in the opposite direction across the saddle (**Fig. 2E**). The glycoprotein-related genes *Edil3*, reported to have elevated expression in the APOE4 genotype,^43^ and *Mgat4c*, identified as a glioblastoma signature gene,^44^ varied more gradually along the path (**Fig. 2F**

#### scDMGC: Morse cell-type taxonomy and cell type annotation

Single-cell and single-nucleus sequencing have enabled increasingly detailed transcriptomic taxonomies of the brain. ^1,4,16,34,38^ Existing methods identify statistically coherent cell populations through hierarchical clustering, graph community detection, machine learning, and related approaches.^11,12,45^ A common workflow selects informative genes and representative cells, constructs a graph of neighborhood relationships, partitions the graph into communities, merges insufficiently separable groups, and recursively repeats this procedure to generate a multiresolution taxonomy.^1–3^

scDMGC provides a complementary framework for simultaneously measuring the intrinsic robustness of Morse-defined cell types and evaluating the consistency of labels derived from an external taxonomy. To illustrate this approach, we constructed a Morse graph from 57,902 cells in the mouse anterior cingulate area (ACA),^5^ a cortical region implicated in contextual fear responses to predatory threat48. The annotated dataset contained 10,503 GABAergic and 47,399 glutamatergic cells. At threshold *J* = 0.30, the Morse graph contained 384 vertices, including 160 GABAergic and 224 glutamatergic cells, each retaining its external subclass and cluster annotation. These included GABAergic *Sst*, *Sst Chodl*, *Pvalb*, *Sncg*, *Lamp5*, and *Vip* subclass populations; glutamatergic L2/3 IT, L4/5 IT, L5 IT, L5 PT, L5/6 NP, L6, L6 IT, and L6b subclass populations; and region-specific types including L4-RSP-ACA and CA2-IG-FC (**Fig. 3A**). Thus, each Morse-graph vertex can be directly compared with its externally assigned taxonomic label.

**Figure 3.**
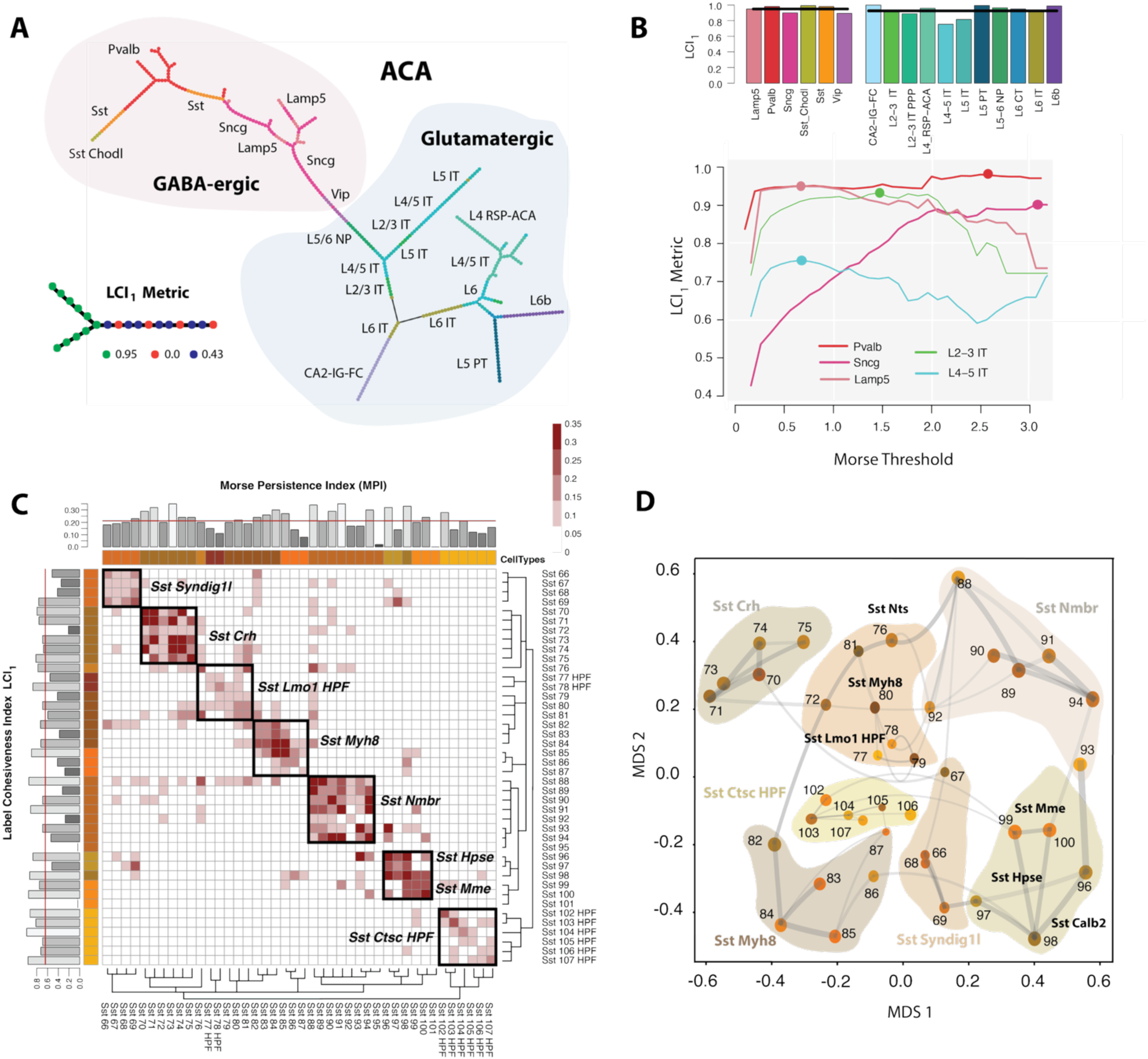
scDMGC Morse cell-type persistence matrix and taxonomy. (A) Morse graph for 57,902 ACA cells^5^ at *J* = 0.30, showing separation of GABAergic and glutamatergic populations and their subclass labels. Inset illustrates the calculation of *LCI*_1_ (**Methods**). (B) *LCI*_1_as a function of persistence threshold for selected GABAergic and glutamatergic subclasses; points indicate the maximum value for each subclass. Bar plot summarizes maximum *LCI*_1_ values across subclasses (**Fig. S4**). (C) Morse Persistence Matrix (MPM). Rows and columns correspond to annotated cell types also identified as Morse cell types. Each matrix entry is the largest persistence threshold at which a minimal Morse-graph connection is retained between the corresponding pair of types. Cluster labels are as defined in Ref. 5. The top annotation shows the Morse Persistence Index (MPI) for each type, defined as the largest *J* at which its representative cell remains a local maximum of the Morse graph. The left annotation shows *LCI*_1_, and color bars indicate supertype membership. Boxes denote Louvain communities (**Methods**) corresponding to supertypes; the dendrogram represents hierarchical clustering from the original annotation.^5^ (D) Constellation-map representation of a Morse Persistence Taxonomy (MPT) obtained by multidimensional scaling of the MPM. Vertex size is proportional to *LCI*_1_, and edge thickness reflects off-diagonal MPM values, which quantify persistent connections between cell types.

The intrinsic robustness of a Morse-defined type is quantified by the *Morse Persistence Index* (MPI; **Methods**), defined for a representative cell as the largest threshold *J* at which that cell remains a local maximum of the Morse graph. Intuitively, MPI is high when a transcriptomic profile is supported by a dense local neighborhood, is well separated from neighboring profiles, or both. When Morse-graph vertices carry labels from an external taxonomy, MPI therefore provides a measure of the extent to which the local transcriptomic structure supports those labeled populations. Analogously, the proximity between pairs of labeled cell types can be quantified from the persistence of Morse-graph connections between representative cells as *J* increases. Collectively, these pairwise relationships define a *Morse Persistence Matrix* (MPM), providing an scDMGC-based representation analogous to cell-type clustering (**Methods**).

A complementary measure evaluates the consistency of external labels along Morse-graph edges. For this purpose, we use the *Label Cohesiveness Index* (LCI), adapted from scan-statistical concepts.^46^ For all Morse-graph vertices carrying label ℓ, *LCI*_1_ is defined as the mean fraction of Morse neighbors carrying the same label (**Fig. 3A; Methods**). Thus, high *LCI*_1_indicates that cells with the same external annotation occupy locally coherent regions of the Morse graph. **Fig. 3B** shows how this coherence varies with persistence threshold for GABAergic and glutamatergic subclasses. Increasing *J* initially removes weakly supported edges and increases label coherence, reaching an optimal range; at higher thresholds, progressively fewer vertices and connections remain, and individual labeled populations eventually lose representation from the graph. At the broadest taxonomic level, GABAergic and glutamatergic vertices were perfectly separated (*LCI*_1_=1.0). Maximum *LCI*_1_ was also high across subclasses (mean 0.949 for GABAergic and 0.924 for glutamatergic populations), supporting the consistency of these labels with the intrinsic structure recovered by scDMGC (**Fig. S4; Suppl. Table 1**).

We used these metrics to analyze 42 brain-wide *Sst* clusters represented in the ACA dataset and defined by the reference taxonomy.^5^ Of the 42 annotated labels, 40 were represented by a Morse maximum, whereas *Sst* 95 and *Sst* 101 were not (**Fig. S5**). The resulting Morse Persistence Matrix (MPM) is shown in **Fig. 3C**. Rows and columns correspond to Morse types labeled according to the reference taxonomy, and each entry gives the largest persistence threshold at which a minimal Morse-graph connection remains between types *i* and *j*. MPI and *LCI*_1_ values are shown as annotation bars in **Fig. 3C**. Mean MPI was 0.21, ranging from 0.00 for *Sst* 101 to 0.35 for *Sst* 91; mean *LCI*_1_was 0.614, ranging from 0.00 for *Sst* 95 to 0.947 for *Sst HPF* 104. Thus, *Sst* 95 and *Sst* 101 showed comparatively weak support for transcriptomic distinctness at the scales examined.

The MPM also recovered coherent supertype-level blocks, including *Sst Crh* (*Sst* 70–75) and *Sst Nmbr* (*Sst* 88–94). Hippocampal *Sst* types had a comparatively low mean MPI of 0.16 but high label coherence (mean *LCI*_1_=0.784), illustrating that persistence and label cohesiveness capture complementary properties of taxonomic organization. Louvain community detection^45^ applied to the MPM identified seven groups corresponding broadly to reference supertypes, whereas off-diagonal entries revealed additional relationships across groups. Multidimensional scaling^47^ of the MPM produced a quantitative constellation map (**Fig. 3D**) that largely recapitulated the reference supertype organization, with notable deviations involving *Sst Calb2*, *Sst Nts*, and *Sst Etv1*. The result is a Morse Persistence Taxonomy (MPT) that generalizes conventional hierarchical organization. An analogous MPM and MPT for glutamatergic L4 IT, L4/5 IT, L5 IT, and L5/6 IT CTX populations is shown in **Fig. S5**.

To extend scDMGC to a larger scale, we constructed an MPM for all 177,594 GABAergic cells.^1,5^ The resulting representation contained 119 Morse representative types and exhibited strong block-diagonal structure across the *Lamp5*, *Pvalb*, *Sncg*, *Sst*, *Sst Chodl*, and *Vip* subclasses (**Fig. 4**). Mean MPI was 0.235 and mean *LCI*_1_ was 0.663. These results are consistent with whole-brain transcriptomic studies showing that major neuronal classes and subclasses are highly reproducible, whereas finer cluster-level distinctions are generally more variable across datasets.^1,4,16,34,38^

**Figure 4.**
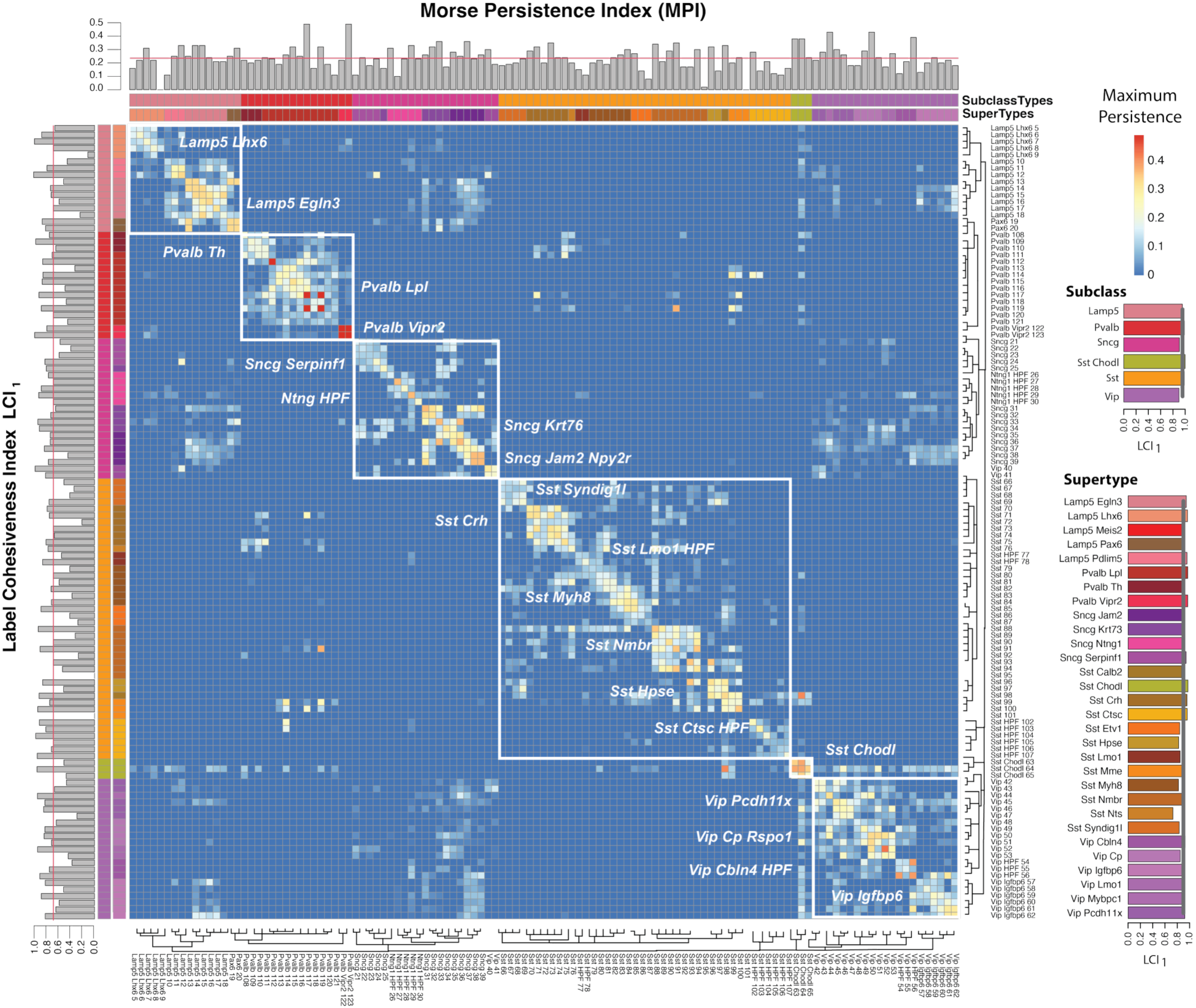
Morse persistence structure of GABAergic cell types. Morse Persistence Matrix (MPM) for 119 Morse representative types from the *Lamp5* (n = 42,144), *Pvalb* (n = 30,461), *Sncg* (n = 13,877), *Sst* (n = 45,467), *Sst Chodl* (n = 1,961), and *Vip* (n = 43,684) subclasses; 21 of 30 reference supertypes are annotated. Matrix entries give the largest persistence threshold at which a Morse-graph connection remains between each pair of types. The top annotation shows the Morse Persistence Index (MPI; mean = 0.235), and the left annotation shows maximum *LCI*_1_ (mean = 0.663). Color bars indicate subclass and supertype membership, and dendrogram clusters correspond to original cell type taxonomy.^5^ Right-hand bar plots summarize maximum *LCI*_1_across taxonomic levels. Label coherence increased with taxonomic level, from clusters (mean maximum *LCI*_1_=0.665) to supertypes (0.923) and subclasses (0.948).

Supertypes provide an intermediate taxonomic level between subclass and cluster and are often more reproducible across datasets.^8,48^ scDMGC quantified label coherence at all three levels directly in the input expression space. Mean maximum *LCI*_1_ increased from 0.665 at the cluster level to 0.923 at the supertype level and 0.948 at the subclass level (**Fig. 4**), indicating that supertype labels were nearly as coherent as subclass labels in the Morse graph. The MPM was broadly consistent with the reference hierarchy (**Fig. 4**, dendrogram) and recovered strongly isolated supertypes, including *Pvalb Vipr2* (*LCI*_1_=0.866, MPI = 0.355), *Lamp5 Lhx6* (*LCI*_1_= 0.647, MPI = 0.182), and *Sst ChodI (LCI*_1_ =0.625, MPI = 0.333), while retaining finer within-supertype variation. The high Morse-graph coherence of supertypes therefore provides quantitative support for this intermediate level of transcriptomic organization between clusters and subclasses.

#### scDMGC and cell-type gradients

Molecular gradients across the cortical sheet, some of which may reflect developmental patterning,^3^ have been described in bulk human transcriptomic data.^35,49^ Single-cell studies similarly show that cell types shared across cortical areas can exhibit graded regional variation.^1,16,38^ More generally, transcriptomic organization can combine discrete cell identities with continuous variation within and between types, ^1,28,5^ raising the question of when cortical populations are best represented as discrete classes versus positions along continuous transcriptional manifolds.

scDMGC provides a direct way to characterize such gradients at cellular-level resolution. A study of 3,663 hippocampal CA1 inhibitory neurons^28^ identified 10 major GABAergic groups comprising 49 fine-scale clusters, while also revealing continuous variation both within and between classes. **Fig. 5A** reproduces a differential-expression-based measure of cluster separation from this dataset. We constructed a Morse graph from the same cells and derived a *Morse Persistence Matrix* (**Fig. 5B**), in which each entry gives the largest persistence threshold at which a graph connection remains between cells carrying the corresponding cluster labels (Methods). The broad structure of the two representations is highly similar (Pearson (vectorized) 0.851), indicating that scDMGC recovers established relationships among CA1 inhibitory cell types directly from local transcriptomic neighborhood structure.

**Figure 5.**
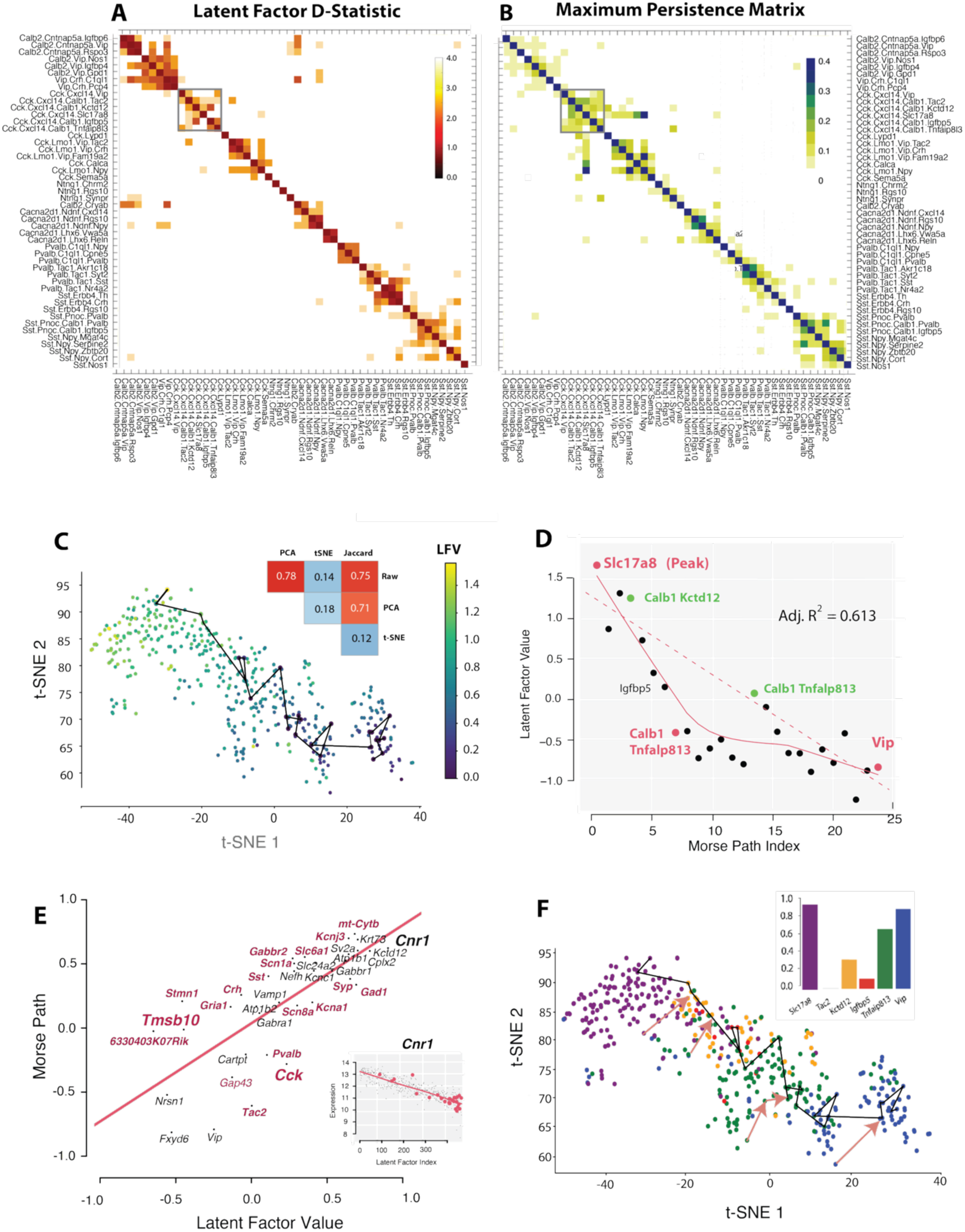
scDMGC resolves a transcriptomic continuum. (A) Reproduction of transcriptomic clusters identified in a study of 3,663 hippocampal CA1 inhibitory neurons.28 The map shows Cohen’s *d* statistic of pairwise cluster separation, with lighter values indicating greater separation and darker values indicating greater mixing.28 (B) scDMGC Morse Persistence Matrix (MPM) for the same populations, showing a similar organization. The box indicates the *Cck Cxcl14*-expressing subcluster spanning the previously identified transcriptional gradient. (C) t-SNE representation of 465 *Cck Cxcl14*-expressing CA1 interneurons,28 colored by latent-factor value, with the 26-cell Morse path containing three peaks and two saddles overlaid. Inset compares correlations between pairwise distances in each representation and pairwise distances in the original input space; (27,998 genes), PCA space (100 dimensions), t-SNE space, and Jaccard space. (D) Latent-factor value as a function of Morse-path position; peaks are shown in red and saddles in green. The linear fit has adjusted (R² = 0.615); curved line Lowess fit (**Methods**.) (E) Correlation of axon-targeting gene expression with latent-factor value versus Morse-path position (Spearman ρ = 0.828; adjusted R² = 0.554, *p* < 1.18 × 10^−7^). Inset shows *Cnr1* expression, with cells on the Morse path indicated in red. (F) Cck expression cells colored by Morse gradient flow assignment. Inset shows agreement of original type with Morse assignment.

Harris et al.^28^ identified a population of 465 *Cck Cxcl14*-expressing cells comprising the *Slc17a8*, *Calb1 Tac2*, *Calb1 Kctd12*, *Calb1 Igfbp5*, *Calb1 Tnfaip8l3*, and *Vip* subclusters that exhibited a continuous transcriptional gradient. The original study represented this gradient using a single latent factor (LFV) fitted by maximum likelihood (color bar, **Fig. 5C**).At a persistence threshold of *J* = 0.20, scDMGC retained a 26-cell Morse path spanning this continuum, with three peaks separated by two saddles. Jaccard-based distances preserved local neighborhood structure more strongly than t-SNE distances and comparably to distances measured in the input and PCA-reduced spaces (**Fig. 5C**, inset).

scDMGC therefore provides a cell-resolved representation of the same transcriptional continuum. The three principal maxima corresponded to representative cells labeled *Slc17a8*, *Calb1 Tnfaip8l3*, and *Vip*. Morse-path position varied nonlinearly with the latent-factor value (adjusted *R*^2^ = 0.61, *p* = 0.074). (**Fig. 5D**), showing that the graph path captures the overall gradient without assuming a linear parameterization. Along the path, mean nonzero expression increased from *Slc17a8* through *Tnfaip8l3* to *Vip*, whereas expression variance and the number of detected genes decreased (**Fig. S6**). The Morse path thus resolves a coordinated transcriptional transition across individual cells while retaining the discrete peaks embedded within the continuum.

Harris et al.^28^ identified 40 axon-targeting genes whose expression varied along the latent continuum. Gene correlations with Morse-path position closely agreed with correlations to the latent factor Spearman ρ = 0.828, adjusted R² = 0.554, *p* < 1.18 *x* 10^−7^; **Fig. 5E**). The cannabinoid receptor *CNR1* (CB1), which mediates endocannabinoid-dependent retrograde regulation of inhibitory and excitatory synaptic inputs onto CA1 pyramidal neurons,^49^ showed the strongest gradient, followed by *mt-Cytb*, *Krt73*, *Kctd12*, and *Gabbr1*. Genes associated with fast-spiking physiology, including *Kcnc1*, *Kcna1*, *Scn1a*, and *Scn8a*, were among those most positively correlated with Morse-path position, consistent with the original analysis.^28^ *Tmsb10* and the neuropeptide genes *Vip*, *Tac2*, and *Cck* showed larger differences between Morse-path and latent-factor rankings (**Fig. S6; Suppl. Table 2**).

A useful feature of scDMGC is that every cell maps through the discrete gradient field to its Morse representative, providing a well-defined association between each cell and transcriptional type or state. Replacing each cell by the expression profile of its Morse representative preserved the dominant transcriptional gradient with little loss of information relative to the original cell-level gradient (Pearson ρ = 0.951, *p* < 8.16 *x* 10^−14^); **Fig. 5F**). However, because the intermediate *Calb1*subtypes *Kctd12*, *Igfbp5*, and *Tnfaip8l3* contributed weaklier to the dominant gradient, assigning each cell the label of its Morse representative showed only partial agreement (49.8%) with the original subtype annotations (inset, **Fig. 5F**). The small *Calb1 Tac2* population (n = 36) was not recovered as a Morse maximum at (J=0.20).

To determine whether these less prominent populations nevertheless corresponded to distinct local transcriptional structure, we lowered the persistence threshold to (J=0.11). At this finer resolution, *Slc17a8*, *Calb1 Tac2*, *Calb1 Kctd12*, *Calb1 Igfbp5*, *Calb1 Tnfaip8l3*, and *Vip* were all recovered as Morse maxima (**Fig. S7**), although the expanded Morse graph corresponded less strongly to the dominant transcriptional gradient. Agreement between Morse-representative labels and the original subtype annotations increased to 70.5% overall (*Slc17a8*, 0.992; *Vip*, 0.833; *Tnfaip8l3*, 0.618; *Igfbp5*, 0.597; *Kctd12*, 0.437; *Tac2*, 0.361). Thus, all six annotated populations correspond to detectable local transcriptional structure, but their prominence depends on persistence scale: only a subset defines the dominant continuum, whereas the remaining populations emerge as distinct maxima at finer resolution. This multiscale result captures the dominant structure at J=0.20 while lower the threshold reveals finer types.

#### Cell-type loss and transcriptional change in Alzheimer’s disease

Alzheimer’s disease (AD), the leading cause of dementia in older adults, is characterized by progressive accumulation of amyloid and tau pathology and profound cognitive decline.^50,51^ The SEA-AD study combined single-nucleus transcriptomics with a BRAIN Initiative reference taxonomy^8,29^ to analyze middle temporal gyrus samples from 84 donors spanning the AD pathological continuum and extending this previously derived taxonomy to 139 supertypes. Quantitative neuropathology also defined a continuous disease pseudo-progression score (CPS) and revealed two broad phases: an early phase marked by slowly increasing pathology, inflammatory microglial states, and loss of somatostatin-positive interneurons, followed by a later phase with rapidly increasing pathology and loss of excitatory neurons and *Pvalb*- and *Vip*-positive interneurons.^8^

The SEA-AD study investigated cell-type vulnerability as a function of CPS, a proxy for disease severity. SEA-AD identified early shifts in *Sst* and *Pvalb* subclass composition, although importantly an apparent reduction in a cell type can reflect either outright cell loss or disease-induced transcriptional changes that impair mapping to the reference taxonomy. To distinguish these possibilities, we constructed a separate Morse graph for each of 109 neuronal supertypes, retaining labels that distinguish cells from early (CPS ≤ 0.5) and late (CPS > 0.5) disease stages. Here, the Morse graphs capture transcriptomic similarity among cells, while the labels record disease state in the SEA-AD data.

For each supertype, we calculated *LCI*_1_ of those cells across persistence thresholds to identify the maximum separability of early and late disease states based on gene expression (**Methods**). **Fig. 6A** then plots these values versus cell type loss as measured by CPS score for all 109 supertypes. Median based quadrants characterize four major categories of cell loss versus transcriptional modification. Across GABAergic and glutamatergic supertypes, the magnitude of cell loss was not linearly related to early-versus-late transcriptional separation (adjusted *R*^2^ = 0.02; **Fig. 6A**); nevertheless, there are distinct patterns that are clear when viewed at the subclass level (**Fig. 6B**). The quadrant representing both substantial cell loss and strong transcriptional change (upper left) was enriched for L2/3 IT supertypes. In contrast, supertypes showing substantial loss but weaker transcriptional separation (lower left) comprised only GABAergic *Sst*, *Pvalb*, and *Sncg* populations, a pattern more consistent with cell depletion than with extensive disease-associated state displacement. A soft-margin linear support-vector machine separated GABAergic from glutamatergic supertypes based on expression with an F1 score of 0.862 (**Fig. 6C**). Although glutamatergic populations contained more variably expressed genes, this difference did not fully account for the CPS effect sizes (**Fig. S8**).

**Figure 6.**
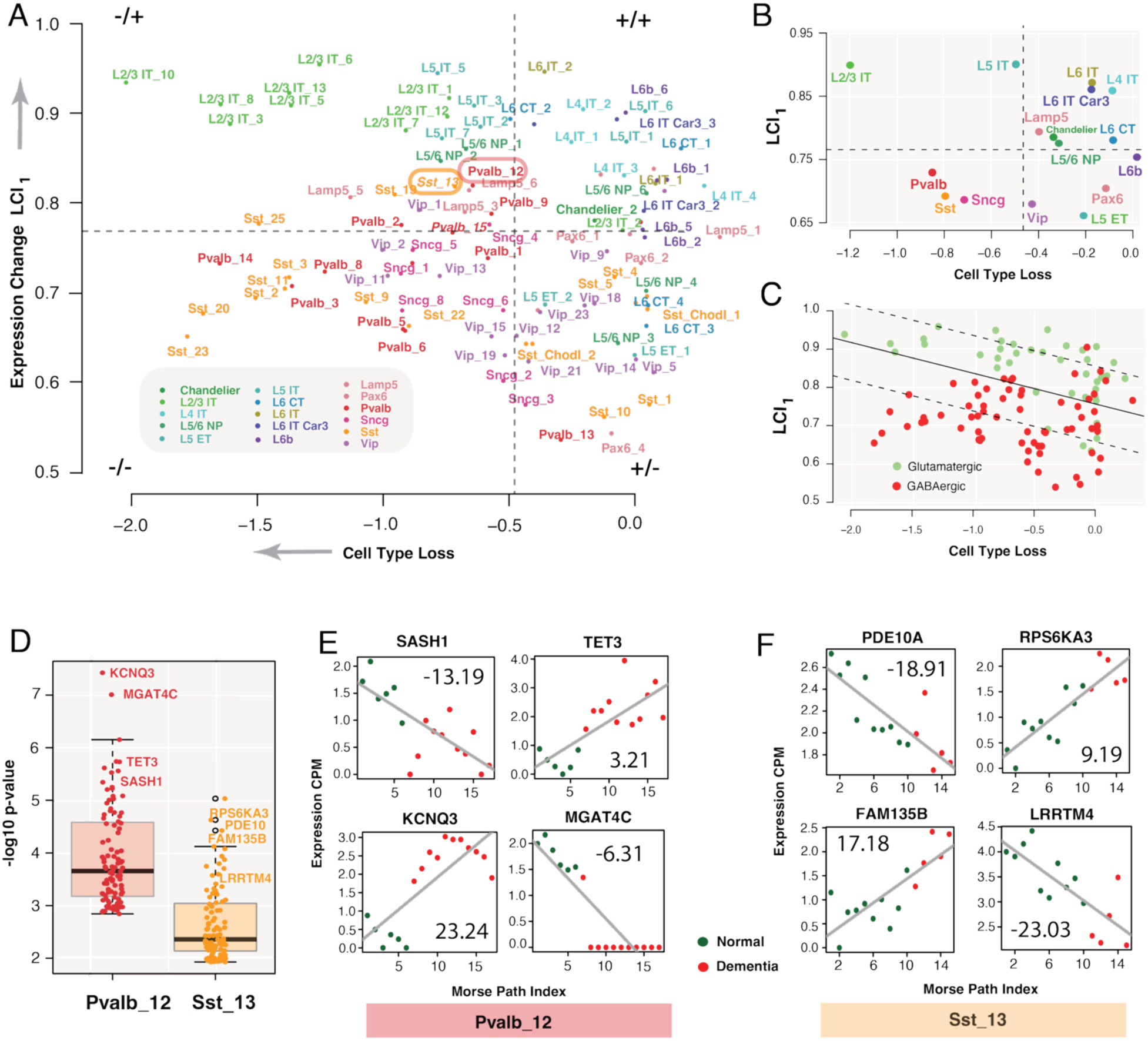
Cell-type loss and transcriptional change in Alzheimer’s disease. (A) Relationship between CPS-associated cell loss (SEA-AD study) (x axis) and early-versus-late transcriptional separation measured by *LCI*_1_ (y axis) across 109 SEA-AD neuronal supertypes.8,29 Dashed lines indicate medians; *Sst* 13 and *Pvalb* 12 are highlighted as GABA-ergic supertypes with moderate vulnerability and high transcriptional variation. (B) Data projected to subclass-level view. (C) Soft-margin linear support-vector machine separating GABAergic and glutamatergic supertypes (F1 = 0.862). (D) Distribution of differential-expression p values for the 100 genes most strongly altered along the *Pvalb* 12 and *Sst* 13 Morse paths. (E) *Pvalb* 12 expression trajectories for genes *SASH1*, *TET3*, *KCNQ3*, and *MGAT4C*; values denote differential effect sizes. (F) *Sst* 13 trajectories for *PDE10A*, *RPS6KA3*, *FAM135B*, and *LRRTM4*.

*Sst*- and *Pvalb*-interneuron dysfunction has been associated with memory impairment in model systems^52,53^ and human AD^54,55^, as well as with disease progression.^56^ We focused on *Sst* 13 and *Pvalb* 12, the GABAergic supertypes with the largest early-versus-late *LCI*_1_ differences (circled, **Fig. 6A**). Morse paths in these supertypes with high *LCI*_1_ showed strong separation of early- and late-disease labels. Differential expression along these paths identified genes with the strongest disease-associated transcriptional transitions (**Fig. 6D**; **Suppl. Table 3**). The effect size was larger for *Pvalb* 12 (Cohen’s D = 1.12) than for *Sst* 13 (D = 0.899), and the *Pvalb* 12 gene set was enriched for a neurofibrillary-tangle measurement term (*p* < 2.59 *x* 10^−9^).^57^

Genes showing the strongest path wise gradients included several previously associated with AD or cognitive phenotype^58,59^ (**Fig. 6E,F**). In *Pvalb 12*, *KCNQ3* increased along the gradient path KCNQ3 contributes to hippocampal spatial coding,^60^ and pharmacological activation of KCNQ/Kv7 channels has been proposed as a potential therapeutic strategy in AD models.^61,62^ In *Sst* 13, *PDE10A* and related phosphodiesterase pathways have been implicated in neuronal signaling and cognitive function, ^63,64^ whereas *RPS6KA3* has been included in a plasma-protein classifier of amyloid burden.^65^ Together, these analyses illustrate how scDMGC can localize disease-associated transcriptional changes to specific paths within defined cell populations. Functional experiments will be required to determine whether these genes are causal contributors to disease progression or potential therapeutic targets.

## Discussion

Single-cell genomics now combines improved molecular sensitivity, multiomic measurements, and increasingly sophisticated computational analysis.^66^ Yet the geometry of omics data remains difficult to characterize: the measurements occupy a high-dimensional input space, the dimensionality and topology of the underlying biological manifold are unknown,^5,31^ and only a subset of expressed genes may be essential for defining cell identity.^17,66^ These uncertainties complicate the identification of stable cell types, continuous variation, and disease-associated states.^67^

Brain cell-type organization is partly hierarchical. Neurons and glia separate at broad levels; neurons divide into excitatory and inhibitory classes; and many subclasses correspond to established anatomical, developmental, and functional properties.^1,16^ At finer resolutions, however, permanent identity can be difficult to distinguish from transient state,^17^ and variation within a type may reflect spatial gradients, activity, development, disease, or transitions between incompletely separated populations.^38,67^ Molecular classifications therefore benefit from independent anatomical, physiological, and functional evidence.^68^ scDMGC addresses one part of this problem by representing both locally distinct transcriptional peaks and continuous paths between them.

The scDMGC method applies discrete Morse theory and persistence to a Jaccard-index field defined on a *k*-nearest-neighbor simplicial complex.^22^ In classical Morse theory, critical points and gradient flows reveal the topology of a manifold;^22^ Forman’s discrete formulation provides combinatorial analogues that can be computed on finite complexes.^22^ In scDMGC, locally dense transcriptional neighborhoods form peaks, persistent drops in similarity from saddles, and optimal gradient paths connect neighboring peaks. The resulting graph is not a conventional low-dimensional embedding: it is a subgraph of the original neighborhood complex and therefore retains local adjacency in the selected gene-expression space.

Across the analyzed datasets, the Morse graphs provided compact summaries of local structure and exposed gradual gene-expression changes that can be visually compressed or fragmented in two-dimensional embeddings.^31^ Genes often changed smoothly as paths approached a saddle, although individual genes could switch on or off near the transition. This representation is well suited to anatomically or transcriptionally proximal populations that exhibit graded variation.^5,16^ It also distinguishes two related questions: whether a labeled population forms a persistent local maximum and whether the label is coherent among neighboring graph vertices.

Trajectory and pseudo time methods also model continuous single cell variation,^67,69,70^ typically by estimating latent variables that order cells along a biological process. scDMGC has a different emphasis. It simultaneously identifies persistent peaks, quantifies the saddles separating them, and reconstructs paths in the observed neighborhood complex. A Morse path should not automatically be interpreted as developmental time, lineage, or causal progression; rather, it represents an optimal route through transcriptionally adjacent cells under the selected metric.

Label evaluation is a particularly useful application. MTM quantifies how persistently a candidate type remains a local maximum, whereas *k*-LCI quantifies the coherence of an external label on the graph. In the *Sst* analysis, these measures supported most reference clusters but provided little evidence for *Sst* 95 and *Sst* 101 at the analyzed scales (**Fig. 4**). The same framework can compare taxonomies across regions, donors, species, or modalities. Because the method requires only a cell-by-feature representation and a distance metric, it could also be applied to chromatin-accessibility data^71^ or used to assess local alignment in multimodal datasets.

scDMGC has several limitations. First, the inferred Morse structure depends on how the transcriptomic data are represented. Normalization, feature selection, the distance metric, neighborhood size (k), sampling density, and the persistence threshold can all influence the locations and prominence of peaks and saddles. These features should therefore be interpreted as structure supported by the chosen representation and scale of analysis, rather than as parameter-independent biological entities. Sparse or uneven sampling can also alter this structure: missing intermediate cells may artificially separate populations, whereas dense sampling of transitional cells may reduce an apparent separation. Second, Morse paths represent routes through transcriptionally adjacent cells and should not, by themselves, be interpreted as developmental trajectories, lineage relationships, or causal mechanisms. Their biological meaning must instead be established using independent experimental or biological evidence.

Finally, computational scalability remains an important limitation. Identifying persistent topological features requires reduction of the filtration boundary matrix, whose dimensions depend on the number of simplices—vertices, edges, and triangles—rather than simply on the number of cells. Classical persistence reduction has worst-case complexity cubic in the size of the filtration, although practical performance is often substantially better because the matrices are sparse and efficient reduction algorithms exploit this structure. ^27^ scDMGC is therefore not intended to replace highly scalable clustering methods for million-cell atlases. Its current strength is the focused analysis of local cell-type structure, transcriptional gradients, and label coherence. Sparse filtrations^72,73^ and distributed or otherwise optimized persistence algorithms^30^ may extend the approach to substantially larger datasets. Within these constraints, scDMGC provides a complementary, metrically grounded representation of discrete and continuous organization in single-cell transcriptomic data.

## Methods

### Mouse hippocampus CA1 dataset

We analyzed 465 *Cck*-expressing inhibitory neurons from mouse hippocampal CA1.^28^ Raw counts for 27,998 genes were transformed as log(CPM + 1). The neighborhood size was set to (k=15) for construction of all (k-nearest-neighbor) k-NN complexes. Additional details of tissue collection, sequencing, and preprocessing are provided in Harris *et al*.^28^

### Mouse cortex and hippocampal formation dataset

We analyzed cells from the 1.3-million-cell adult mouse isocortex and hippocampal formation dataset.^5^ Expression values were transformed as *log*(*CPM* + 1). For each analysis, the 4,000 most variable genes were selected by ANOVA. All k-NN complexes were constructed using (k=25). Additional dataset and preprocessing details are provided in Yao *et al*.^5^

### Human SEA-AD dataset

We analyzed 992,281 GABAergic and glutamatergic neurons assigned to 109 of the 139 SEA-AD supertypes.^8^ Within each supertype, expression values were transformed as [transformation], and the 2,000 most variable genes were retained. All k-NN complexes were constructed using (k=15). For comparisons between early- and late-stage specimen histology, label cohesiveness was evaluated separately for the two histological classes at each persistence threshold. The reported *LCI*_1_ score was the maximum, across thresholds, of the smaller of the two class-specific *LCI*_1_ values, thereby requiring both classes to exhibit local label coherence. Dementia-associated gene-expression changes were evaluated along Morse paths connecting cells from donors without dementia to cells from donors with dementia. Additional dataset and donor information is provided in Gabitto *et al*. ^8^

### Topological data analysis

scDMGC uses standard concepts from computational topology and discrete Morse theory.^19,22^ Cells are represented as points in the selected gene-expression space, and Euclidean distance is used to construct a k-nearest-neighbor graph. The directed neighbor relation is converted to an undirected graph where an edge is retained when either cell is among the other cell’s k nearest neighbors (union). A simplicial complex is composed of simplices of increasing dimension: vertices are 0-simplices, edges are 1-simplices, and triangles are 2-simplices. scDMGC uses the two-dimensional simplicial complex induced by the k-NN graph. Each cell corresponds to a vertex, each retained k-NN relationship to an edge, and every three-clique in the graph is filled to form a triangle. No simplices of dimension greater than two are included.

### Jaccard field and lower-star filtration

A filtration is a nested sequence of simplicial complexes in which simplices are added according to an ordering function. scDMGC constructs a simplex-wise lower-star filtration of the two-dimensional k-NN complex using local neighborhood overlap. For an edge *(u,v)*, let *N_k_*(*u*) and *N_k_*(*v*) denote the k-NN neighborhoods of cells *u* and *v*. Their *Jaccard similarity* is defined as the fraction of neighbors shared by the two cells.

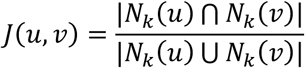

The *Jaccard index* assigned to a vertex *u* is the maximum edge similarity over edges incident to that vertex, *J*(*u*) = *max*_{(*u*,*v*)∈_*_E_ J*(*u*, *v*).

Triangles inherit a filtration value determined by their constituent edges so that the filtration respects the simplicial face relation. Specifically, the value assigned to a triangle is the minimum (or inverse-filtration equivalent) of the values of its three edges. The filtration is ordered by inverse Jaccard similarity so that cells with strongly overlapping neighborhoods enter at higher-density portions of the underlying transcriptional landscape. Because neighborhood overlap takes only a finite set of values, multiple vertices or simplices can have identical filtration values. These ties are resolved deterministically by ordering simplices first by dimension and then lexicographically by their ordered vertex indices, producing a simplex-wise filtration. Thus, faces precede cofaces when they share the same filtration value. This ordering can affect which individual cell is selected as a representative when several cells have identical Jaccard values, but it does not change the underlying tied filtration level.

Relative maxima of the Jaccard field correspond to cells whose local neighborhoods exhibit particularly strong transcriptional similarity. These maxima form candidate peaks in the discrete Morse representation. Persistence measures the prominence of such peaks relative to the saddles through which they connect to neighboring peaks. For further details on filtrations, discrete Morse theory, and persistence computation see.^18,22,27^

### scDMGC algorithm

The original discrete Morse graph-construction algorithm extracts ridge structure from a sampled density field.^27^ It first constructs a lower-star filtration of the discretized domain using the negated density function and then returns the unstable one-manifolds associated with persistent critical points. The generalized algorithm accepts an arbitrary filtration and produces a lexicographically optimal graph with respect to that filtration. For scRNA-seq data, we replace the weighted Rips filtration used in the point-cloud implementation with the Jaccard-ordered *k*-nearest-neighbor filtration described above. The parameter δ controls persistence-based simplification: larger values retain only more prominent graph features. In this study, δ is expressed in units of Jaccard similarity and is denoted *J* in the Results.

**Algorithm: scDMGC (*X*, *k*, *δ*)**

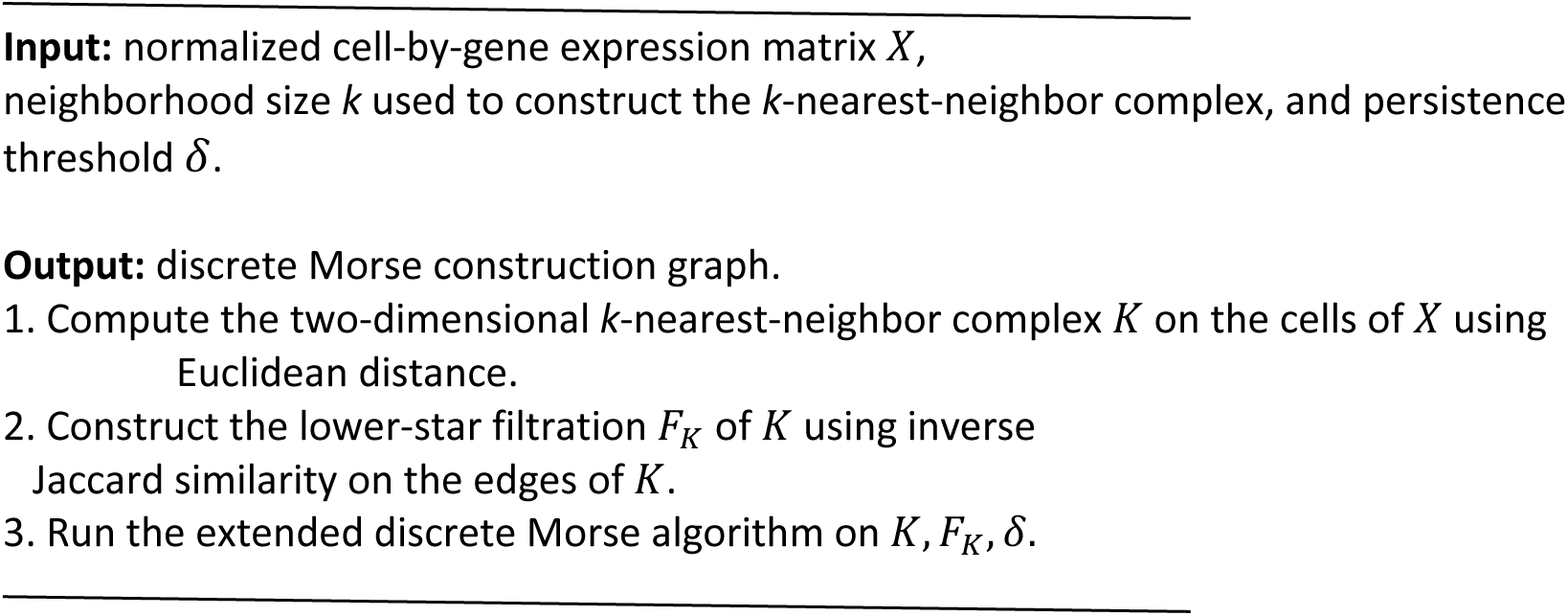

### Wasserstein distance

Cell distributions were compared using the Wasserstein distance,^36^ implemented with the Python Optimal Transport (POT) library.^74^

### Weisfeiler–Lehman graph distance

Morse graphs were compared using the Weisfeiler–Lehman graph distance,^75^ an attributed-graph distance based on iterative Weisfeiler–Lehman neighborhood refinement^37^combined with an optimal-transport comparison of the resulting graph representations.

### Soft-margin support-vector machine

A linear soft-margin support-vector machine in R was used to distinguish GABAergic from glutamatergic supertypes. The classifier was trained using *LCI*_1_versus cell type loss. Classification performance was evaluated by cell type separability.

### Label cohesiveness index (LCI)

Given a graph G(V, E) and a label function *L*: *V* → {*l*_1_, *l*_2_, …, *l_n_*}, the label cohesiveness index *LCI*_1_for label *l_i_* is defined as follows: For every vertex *v* carrying label *l_i_* we identify the fraction of neighbors carrying the same label. These counts are averaged across all vertices carrying *L*. Thus, LCI is the fraction of vertices within graph radius 1 of label vertices that also carry label *l_i_*. The focal vertex itself is excluded. Values near 1 indicate that the label occupies a locally coherent region of the Morse graph, whereas lower values indicate mixing with other labels. The generalization *LCI_k_* to vertices reachable in k steps is straightforward but did not measurably change results.

### Morse-graph separability

To quantify regional separability, we considered all triplets (a, b, c) in which a and b belong to the same class and c belongs to a different class. Separability is the fraction of triplets satisfying *d*(*a*, *b*) < *min* {*d*(*a*, *c*), *d*(*b*, *c*)}. Thus, the measure estimates the probability that two observations from the same region are more like one another than an observation drawn from a different region.

### Morse Persistence Index

For a putative cell type *c*, the Morse Persistence Index (MPI) was defined as the largest persistence threshold J at which the representative of *c* remained a relative maximum of the simplified Morse graph. A large MPI therefore indicates that a type is supported by a locally dense and internally similar transcriptional neighborhood that remains distinct across a broad range of persistence thresholds.

### Morse Persistence Matrix

For N labeled cell types, the Morse Persistence Matrix (MPM) is an *NxN* symmetric matrix describing the persistence of graph connections between pairs of types. Entry *M_i_*_j_ is the largest persistence threshold J at which the simplified Morse graph retains at least three noncritical edges connecting vertices assigned to types *i* and *j*. Requiring multiple connecting edges reduces sensitivity to isolated cross-type edges and was used as an empirically selected robustness criterion for declaring a persistent relationship between two types. For the complete GABAergic analysis, MPMs were first computed separately within the *Lamp5*, *Pvalb*, *Sncg*, *Sst*, *Sst Chodl*, and *Vip* subclasses. Additional pairwise MPMs were then computed for every pair of subclasses. The final combined matrix used the corresponding within-subclass MPM entry for pairs of types belonging to the same subclass and the relevant pairwise-subclass MPM entry for types belonging to different subclasses.

### Constellation-map embedding

Types that were outliers for both maximum 1-LCI and maximum persistence were excluded. We applied multidimensional scaling to the MPT to obtain two-dimensional type coordinates. Vertex size was proportional to maximum 1-LCI across thresholds. Louvain community detection was used to estimate supertype groupings. An edge was drawn between two types when the corresponding MPT entry was at least 0.05 and at least 75% of the maximum entry in either type’s MPT row; edge thickness was proportional to the MPT value.

### Morse Persistence Taxonomy and constellation-map embedding

The MPM was used to construct the Morse Persistence Taxonomy (MPT), which summarizes persistent pairwise relationships among labeled cell types. Multidimensional scaling was applied to the persistence-based type representation to obtain two-dimensional coordinates for the constellation map. Vertex size is proportional to the maximum *LCI*_1_ attained across persistence thresholds, and Louvain community detection^76^ is used to identify groups of transcriptionally related types. Types that were outliers with respect to both maximum *LCI*_1_ and maximum Morse persistence were excluded before visualization. An edge was drawn between types *i* and *j* when their MPM entry was at least 0.05 and at least 75% of the maximum MPM value in the row corresponding to either type. Edge thickness was proportional to the corresponding persistence value.

## Data analysis

Statistical and computational analyses were performed using R version 4.3.1 and Python version 3.12.

## Acknowledgments

The authors thank Allen Institute founder Paul G. Allen for his vision, encouragement, and support. We thank Marcus Hooper of the Allen Institute for providing the initial scRNA-seq datasets used to develop the method.

## Supplemental Information

**Figure S1.**
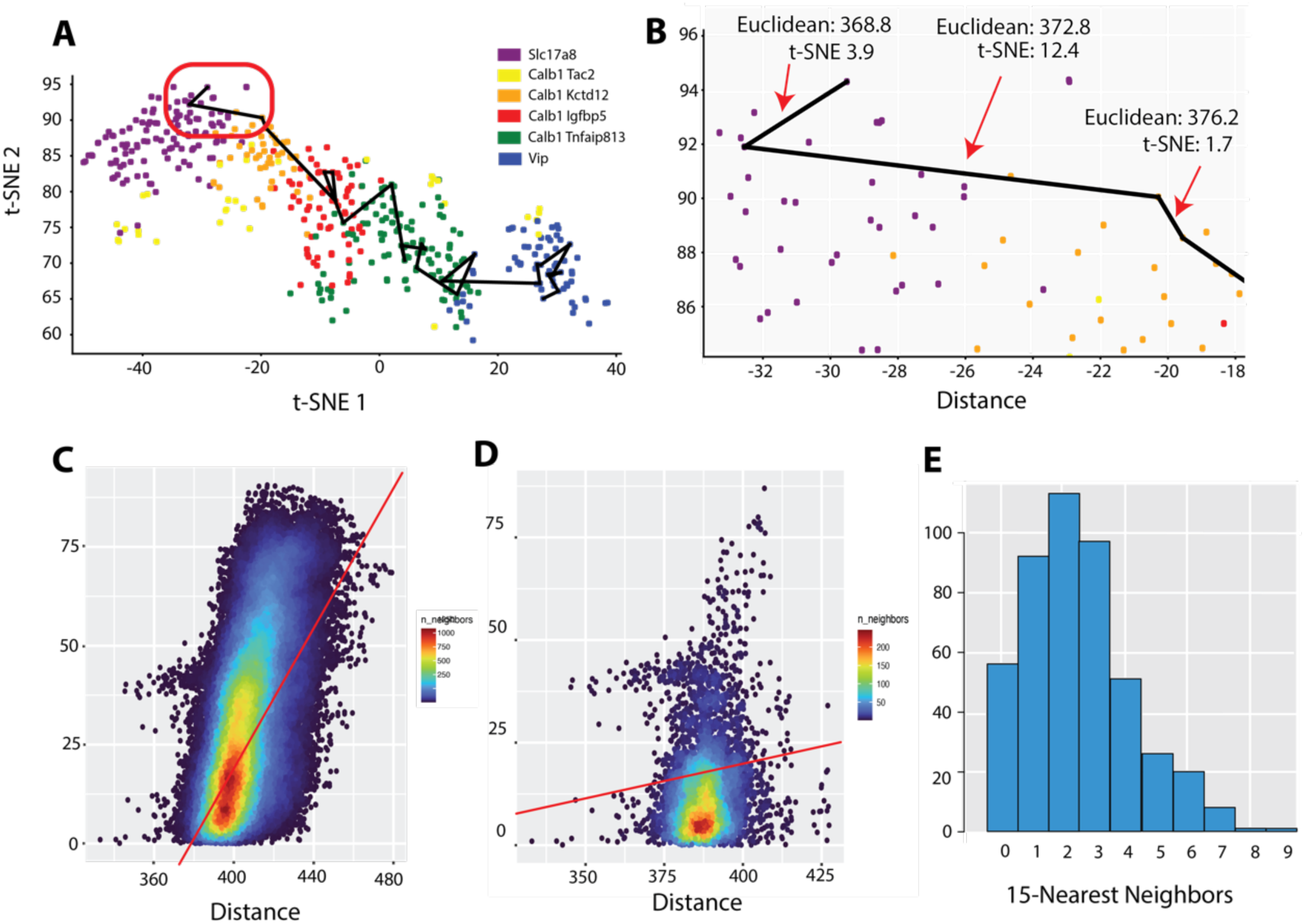
(A) *Cck* cell dataset of 465 cells from^1^ colored by cell type and the Morse graph computed in the raw gene space both embedded in the t-SNE projection. (B) Zoom-in focusing on 3 edges in the Morse graph projected into t-SNE embedding with raw gene space and t-SNE embedding distances. Metric distortion is clearly observed in the embedding. (C) Scatterplot heatmap of the distance between cells in the raw gene space (x-axis) and distance between cells in the t-SNE embedding (y-axis) (⍴=0.6, *p* < 2.2 *x* 10^−16^). (D) The same plot but only including cells that are 15-NN in the raw gene space (⍴=0.1, < 2.2 *x* 10^−16^) showing local structure is not preserved. (E) Histogram of the number of raw gene space 15-NN that are preserved in the t-SNE embedding for each cell. More than half of cells have 2 or fewer nearest neighbors preserved in the embedding.

**Figure S2.**
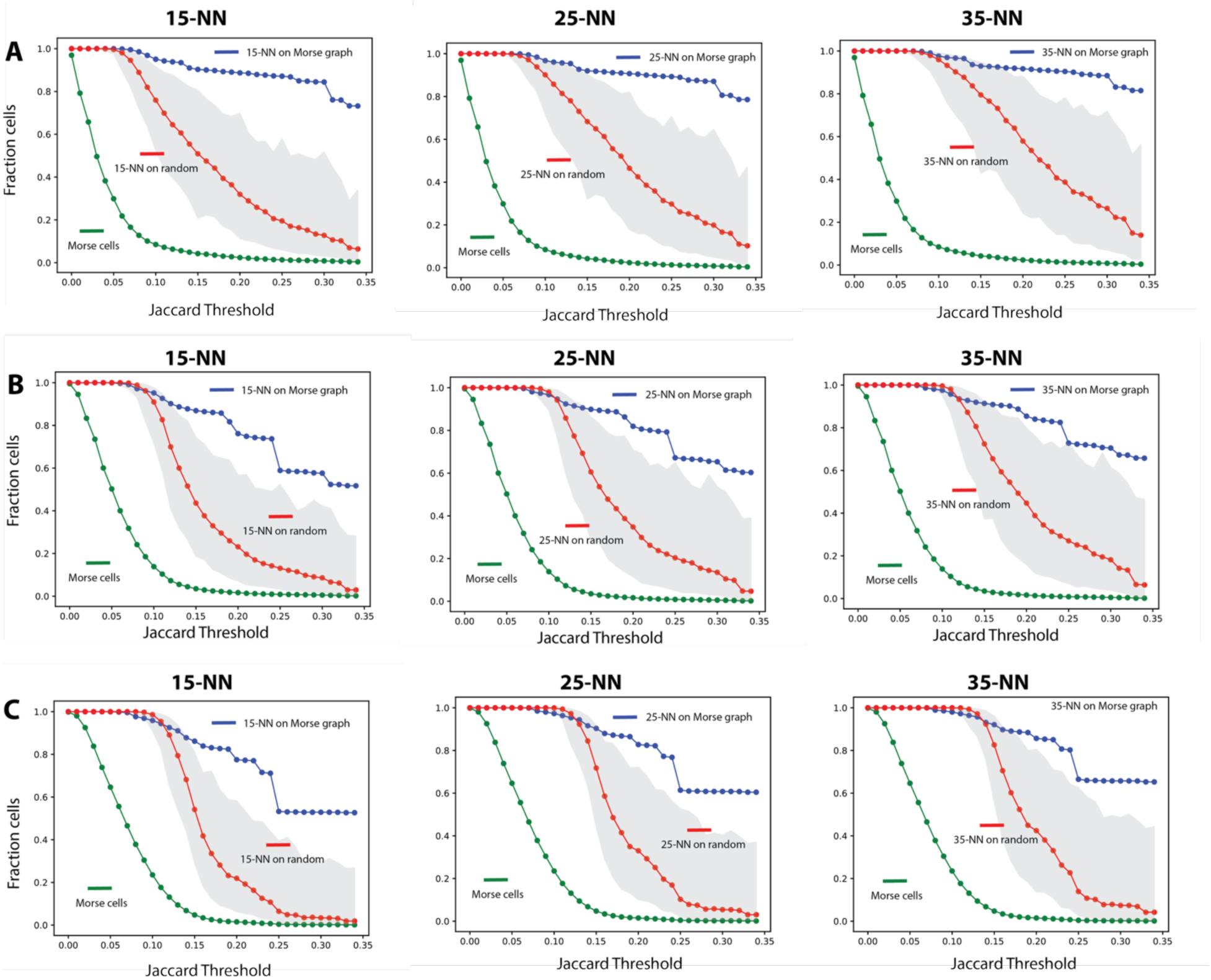
The Morse graph construction is a stable and compact representation for a range of nearest neighbor sizes. Rows (A-C) have been constructed building Morse graphs using 15, 25, 35-NN. Each plot shows the fraction of cells from^2^ on the Morse graph as a function of Jaccard threshold (green curve.) Within each row we examine the structure of the Morse graph over the same range of nearest neighbors. For example, in row B column 1, the Morse graph is constructed using 25-NN. The plot shows the fraction of cells having a 15 nearest neighbor on the graph as a function of threshold (blue curve.) The red curve shows the mean results over 100 random subsets of cells (grey range). At lower thresholds (J) a large fraction of cells remains on the graph (green curves A-C) with sharply decreasing number as threshold increases. Selecting a variable number of nearest neighbors k=15, 25, 35 corroborates the compactness of the Morse representation. A large fraction of neighbors remain on the Morse graph as J increases and remains compact and spanning with respect to random subsets of equivalent size.

**Figure S3.**
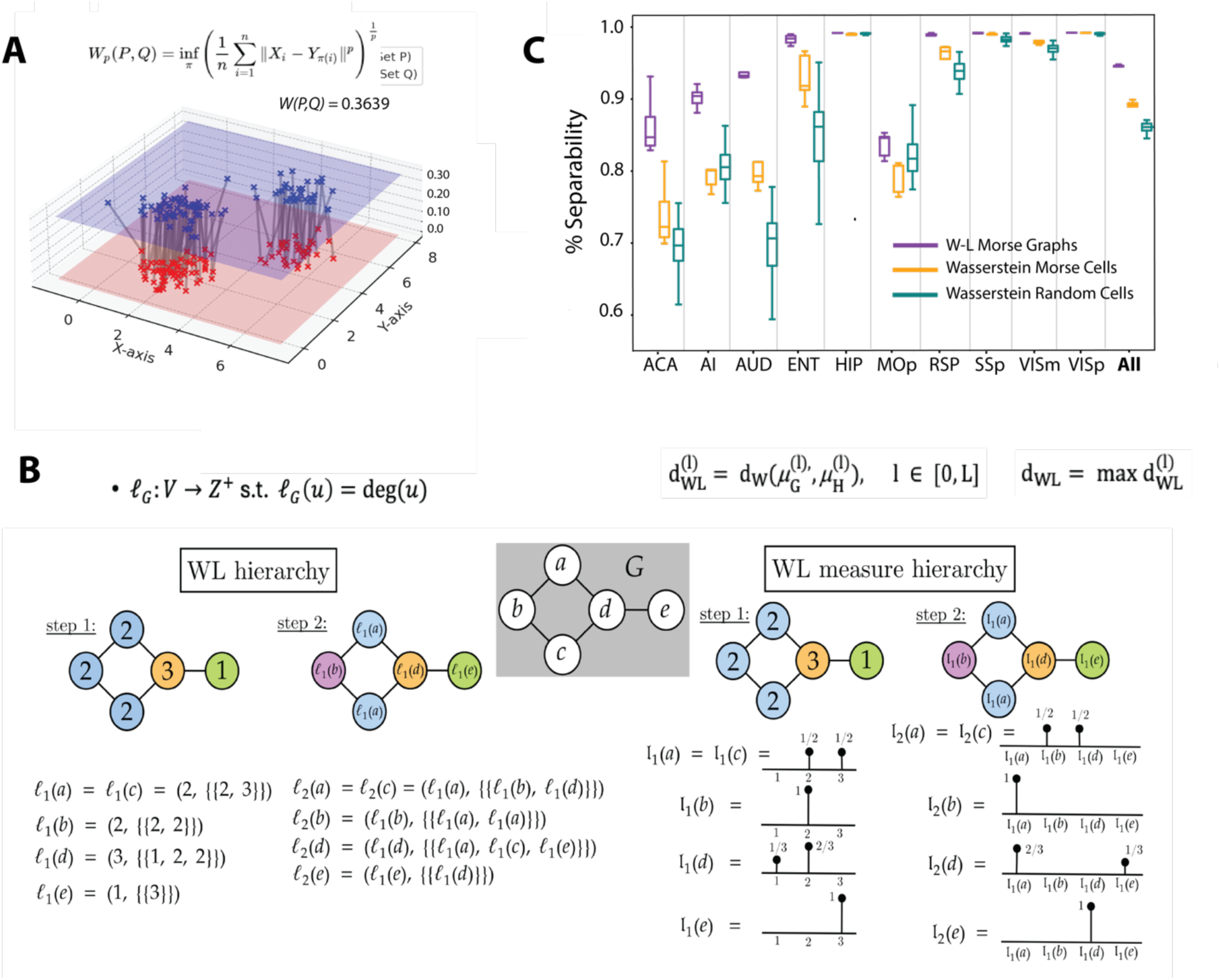
(A) The Wasserstein distance, a special case of optimal transport, is widely used to compare two probability distributions defined on the same space. The *Wasserstein distance* is defined as this minimal transport cost. See^3^ for more precise definitions and details of the Wasserstein distance and related concepts. The Wasserstein distance^3^ *d*_W_(*ρ*, *μ*) *d_w_*(*ρ, μ*) can be used to compare two-point sets *P* and *Q* from *R^d^*, with discrete measure distributions *ρ* and *μ* respectively. Given a set of points *P* = {*p*_1_, …, *p_n_*}, and empirical distribution 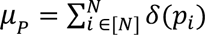 where δ(x) is the Dirac Delta function at *x*, we can then compare the two-point sets *P* and *Q* by the Wasserstein distance between these induced by *μ_P_* and *μ*_Q_. If *P* is a set of points sampled uniformly randomly from a well-behaved continuous measure μ, then it is *d*_W_(*μ_P_*, *μ*_Q_) tends to *0* almost surely as the number of points in *P* goes to infinity. We use the Wasserstein distance to compare the distributions of cells in the high dimensional space of RNA expression data from the Python Optimal Transport library.^4^ (B) The Morse graph contains additional structure in the edges connecting cells. Graph distance measures based on graph edit distance are computationally inefficient even to approximate within a constant factor. An approximate measure can be derived based on the Weisfeiler-Lehman (WL) test, a classical procedure for graph isomorphism testing.^5^ We use the Weisfeiler-Lehman (W-L) distance^6,7^, a notion of distance between labeled measure Markov chains (LMMCs), of which labeled graphs are special cases, and that has been widely used both for designing graph kernels and for analyzing graph neural networks. Computing W-L distance between two graphs G and H amounts to hierarchically computing the measures *μ_G_* and *μ_H_* from graph edge adjacencies and comparing these measures with the Wasserstein distance 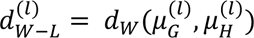. Panel B (reproduced with permission from^6,7^ illustrates computation of the W-L measure for a toy 5 node graph G shown in the middle of the panel. At the zeroth step *l*_5_(*v*) is defined as the degree of vertex v. In the first step, the function *l*_1_(*v*) defined as the set consisting of the degree of v and set of degrees of each of v’s neighbors in G. Each subsequent step iteratively replaces a node with a set adjoining the set of labels from the previous set. The process can be formulated as a Markov chain that explores the connectivity structure of the graph and has appropriate convergence properties. The process is run to convergence defining 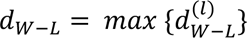. W-L distance captures and compares subtle structures of the underlying LMMCs and is more discriminating than the classical distance measures between graphs. (C) To show that scDMGC Morse graph can differentiate the identity of cortical regions we performed an experiment using *Vip* positive cells from 18 cortical regions including hippocampus.^2^ Ten subsets of 500 cells from each region were used to construct 180 Morse graphs sampling the cell type structure of the 18 distinct regions. To compare cortex region identity based on gene expression we compare distances between sets of cells from different regions using the Wasserstein distance. To compare cortex region identity based on gene expression we compare distances between sets of cells from different regions using the Wasserstein and distances between Morse graphs constructed on sets of cells using W-L distance. For each cortex region *R*, we choose two subsets (A, B) from *R* and the third *C* from a different region. We then measure how close sets of cells and Morse graphs are using these distance metrics. Percent separability is defined as the fraction of triplets for which *W*(*A*, *B*) < *min* {*W*(*A*, *C*), *W*(*B*, *C*)}, indicating that A and B are more similar in transcriptomic structure than either is to C. Panel C shows the results for 10 regions (ACA, AI, AUD, ENT, HIP, MOp, RSP, SSp, VISm, VISp) and where *All* indicates all 18 regions.^2^ Scoring of separability using Wasserstein distance of cells on the Morse graph alone, and W-L graph distance to compare Morse graphs with edge structure. Panel C shows that comparing Morse graphs using the W-L metric captures regional transcriptomic identity significantly better than the W metric on Morse cells alone, as well as the W metric on random cell sets. Over all regions, W-L graph distance attains 0.95 mean accuracy, Wasserstein distance (Morse) 0.88, and Wasserstein distance (random) 0.85, illustrating that Morse graphs capture the regional identity of cell type architecture via cells on the graph, which is further improved using the edges that define their relationships.

**Figure S4.**
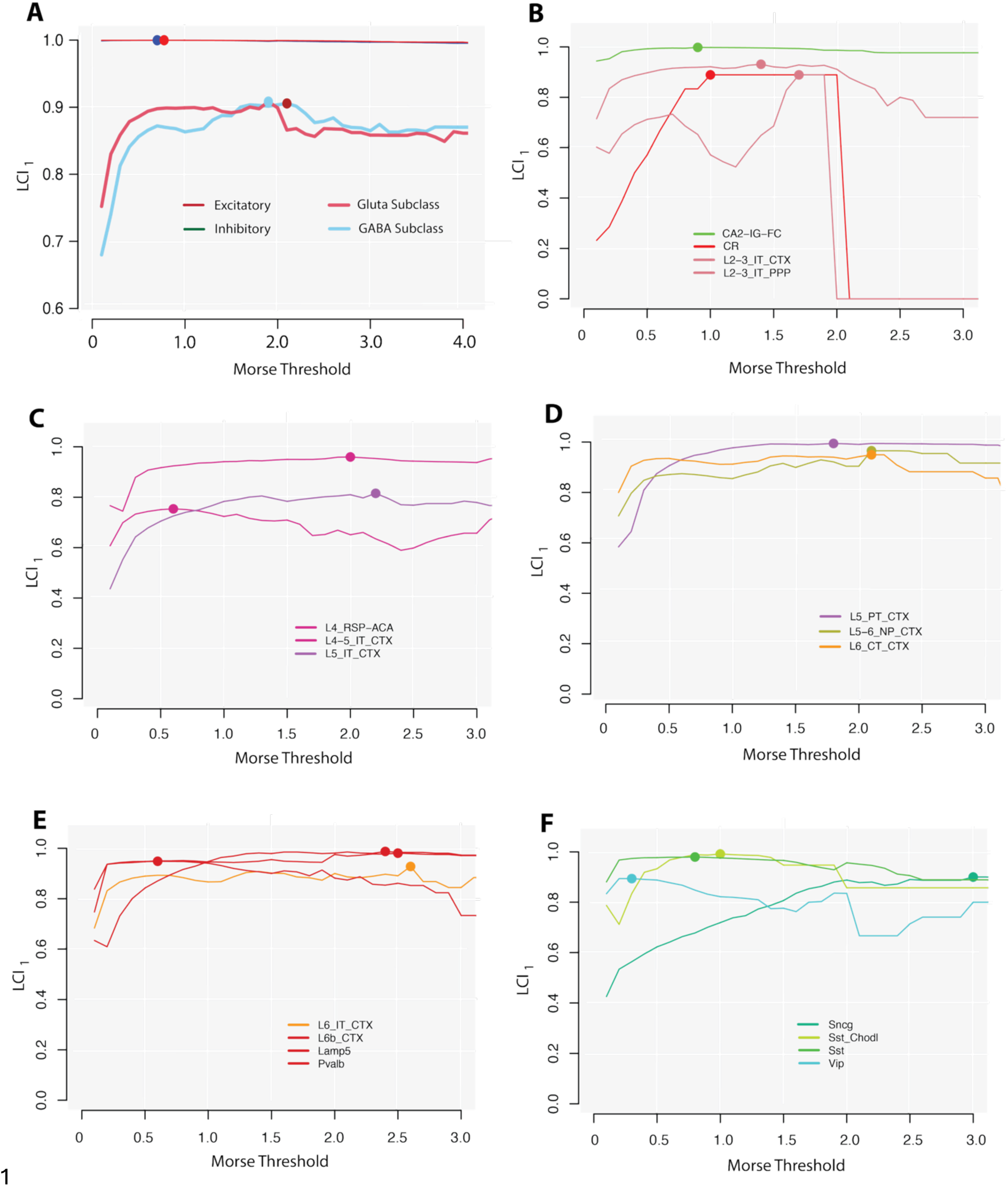
(A) *LCI*_1_metric as a measure of cluster label stability. Highest lines show essentially perfect separation of GABAergic and glutamatergic cell types. Point indicates maximum Morse threshold for maximum *LCI*_1_. Mean of GABAergic and glutamatergic subclasses. (B-F) *LCI*_1_ curves with maxima denoted for GABAergic and glutamatergic subclass types.^2^ *LCI*_1_ depends on the Morse threshold and increases to a point of maximum cell type specificity. The maximum *LCI*_1_value is high for all subclass types (Inhibitory 0.949, Excitatory 0.924) and is consistent with the uniqueness of these cell classes.

**Figure S5.**
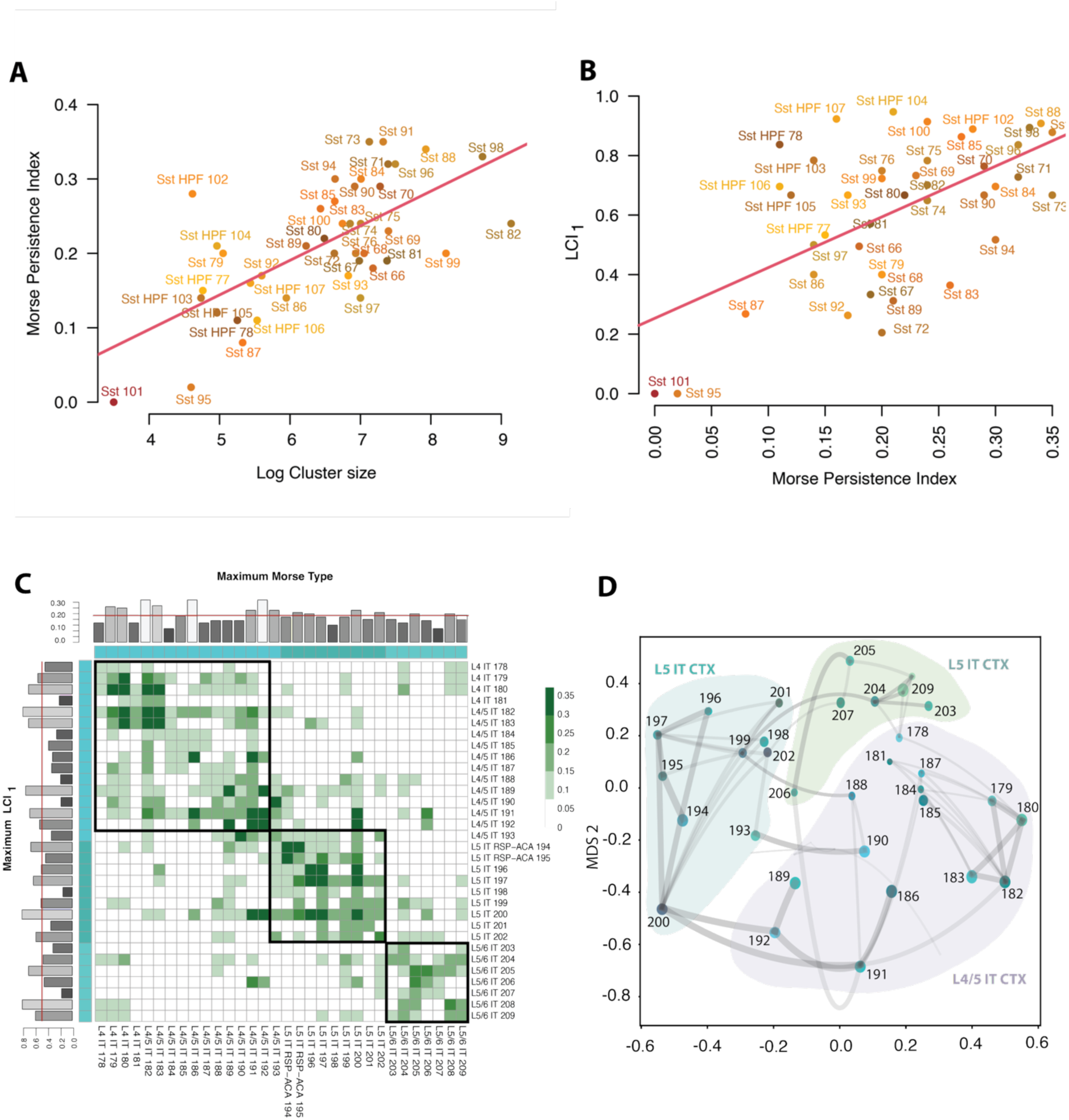
(A) Plot of log of cell type cluster size as identified in^2^ versus Morse Persistence Index (MPI), the largest J for which *c* occurs on *M_J_* as a relative maximum. The measure *MPI*(*c*) increases as the fraction of neighbors of *c* with similar expression increases and is a natural measure of cell type uniqueness. MPI is correlated (⍴=0.666) with the log of cluster size for 42 *Sst* cell types. *Sst 95* and *Sst 101* are removed from the taxonomy. (B) MPI is correlated with *LCI*_1_ label consistency (⍴=0.577) and is measure of agreement or divergence of Morse taxonomy with the annotated labels. (C) The Morse Persistence Matrix (MPM) derived for glutamatergic *L4 IT*, *L4/5 IT*, *L5 IT*, and *L5/6 IT* cortex. Boxed types are three optimal Louvain groups that agree with.^2^ (D) Visual constellation map representation is depicted like Fig. 4D. Morse cell types are input to multidimensional scaling (MDS) to obtain the coordinates for each type. The size of the vertices scales as the maximum *LCI*_1_ score with edge thickness determined by the magnitude of off-diagonal MPT entries and representing cell type confusion.

**Figure S6.**
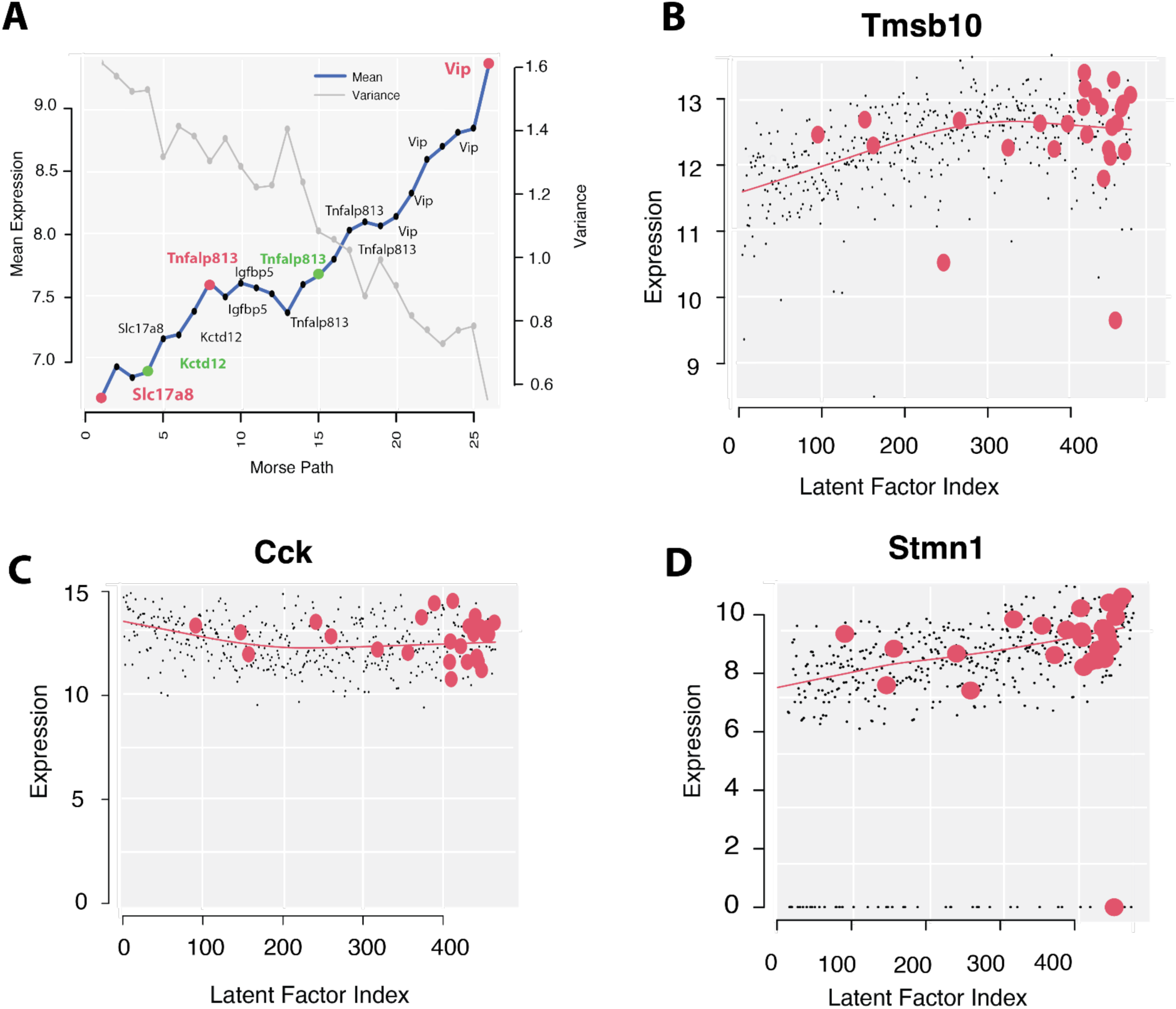
(A) Mean and variance of expression along the Morse path gradient showing peak (red) and saddle cells (green) labeled. The percentage of zero expressing genes is increasing along the gradient (not shown). (B,C,D) Scatter plot with trajectory lowess fit (f=1/3) of expression along latent factor variable (LFV) ordered cells with Morse cells highlighted in red. (B) Thymosin beta 10 (*Tmsb10*), actin processing gene,^8^ (C) Neuropeptide cholecystokinin (*Cck*) soma-targeting basket cells^9^ are expressed ubiquitously in all cells of this data. (D) Stathmin 1 (*Stmn1)*, another actin processing gene. ^10^ All three genes exhibit higher variability on the Morse path. (**Suppl. Table 2**).

**Figure S7.**
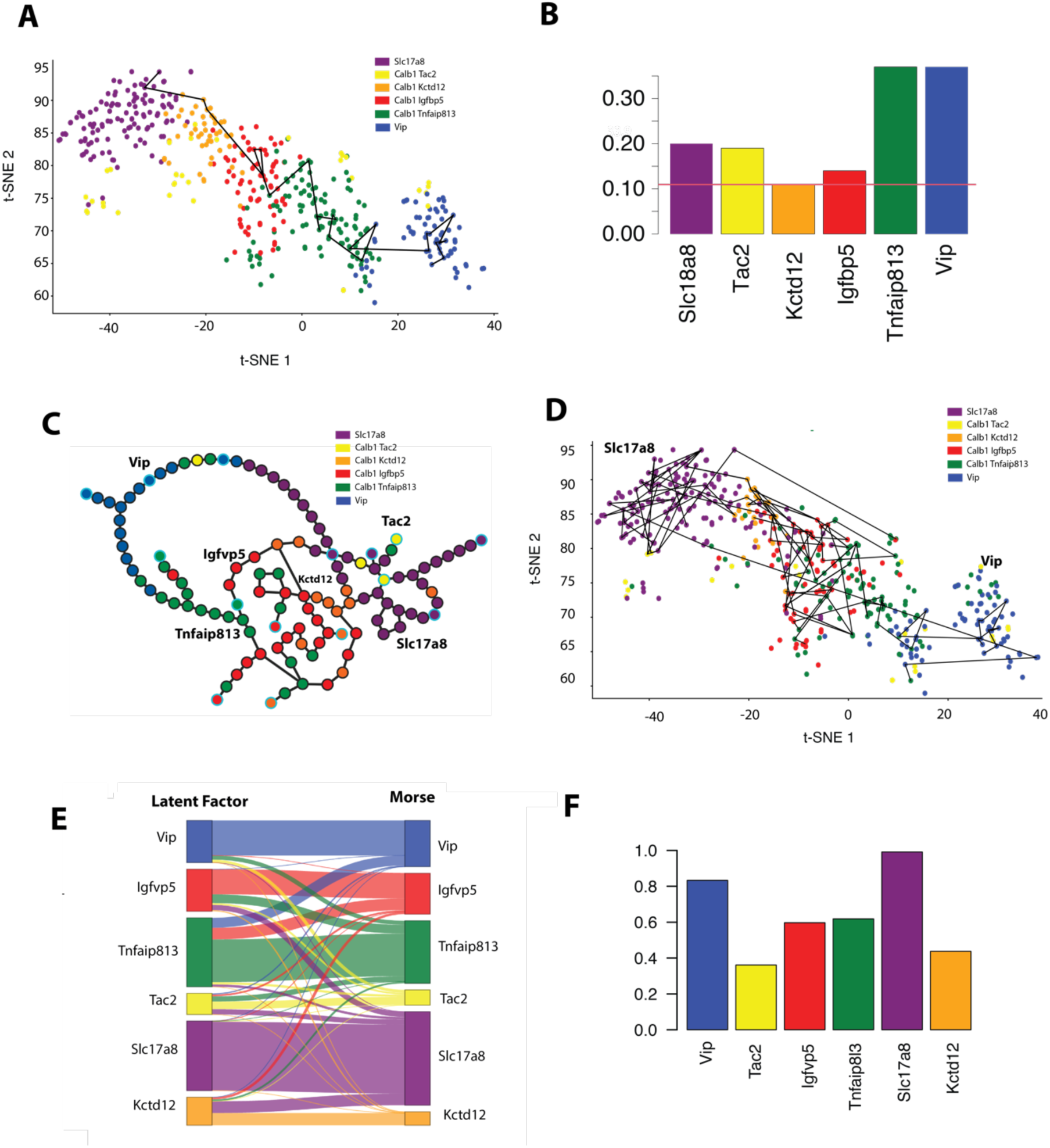
(A) Harris et al^1^ use a modified t-SNE called *tnb-SNE* to visualize the locations of the cells deemed appropriate for data following a negative binomial distribution. 465 cells from the *Cck Cxcl14* class together with labeled cell types *Slc17a8*, *Calb1 Tac2*, *Calb1 Kctd12*, *Calb1 Igfbp5*, *Calb1 Tnfaib813*, and *Vip* and Morse path with L=26 cells. (B) Maximum Jaccard threshold in k-NN graph for which each Harris^1^labeled type occurs. Minimum is J=0.11 for type *Calb1 Kctd12.* (C) Morse graph for the J=0.11 threshold with labeled type from. Maximal Morse types are shown as cyan circled. (D) t-SNE embedded with Morse graph shown and cells colored by gradient flow Morse representative. Every cell in the dataset flows via the underlying gradient path into a single cell on the Morse graph, the Morse representative of the cell. (E) Sankey flow plot showing how latent factor predicted cells are reassigned by Morse gradient flow representatives.(F) Panel shows the original assignment^1^of cells to the types (*Slc17a8* 0.992, *Vip* 0.833, *Tfnfaip813* 0.618, *Igfvp5* 0.597, *Kctd12* 0.437, *Tac2* 0.361). Overall agreement is now 70.5%.

**Figure S8.**
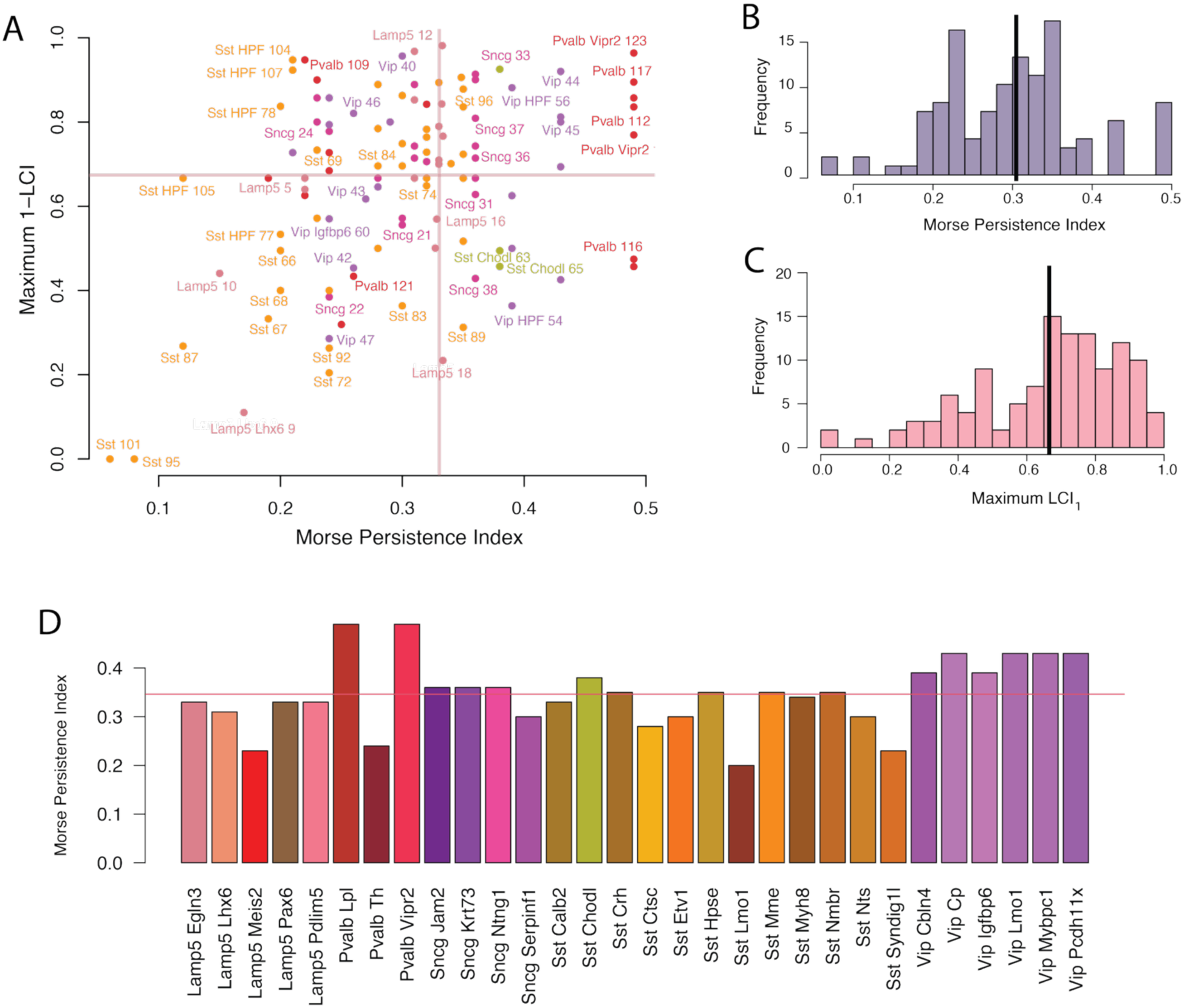
(A) Scatter plot of ACA GABAergic supertypes showing relationship of Morse Persistence Index and maximum *LCI*_1_. Lines indicate mean values in each dimension. Cell type colors as in original publication.^2^ (B) Histogram frequency of Morse threshold for GABAergic types with mean, (C) Maximum *LCI*_1_ distribution. (D) Morse Persistence Index of each ACA GABAergic supertype.

**Figure S9.**
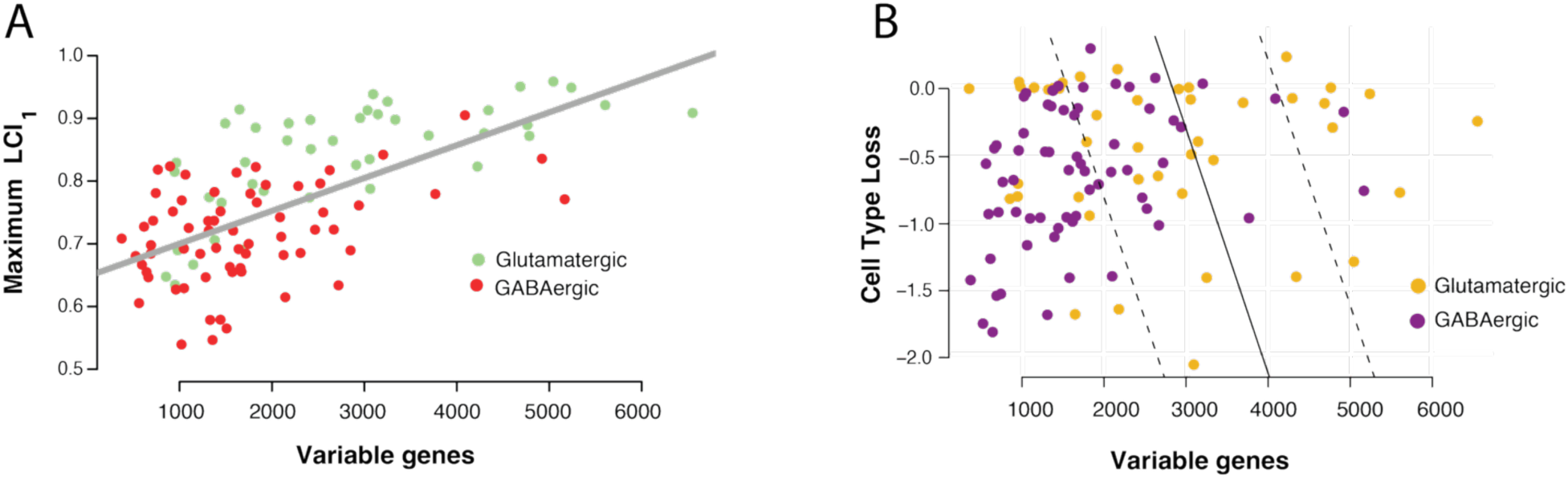
(A) Variability in gene expression was measured in^11^ over profiled cells finding that glutamatergic expressing neurons had a substantially larger number of variable genes. Maximum separability of supertypes based on maximum *LCI*_1_increases by the number of variable genes coded here by GABAergic or glutamatergic expressing supertypes. Adjusted R-squared: 0.3821, p-value: *p* < 4.81 *x* 10^−13^. (B) Effect loss is not fully explained by the increased number of variable genes. Soft margin SVM fitting variable genes to CPS loss effect size separates GABAergic and glutamatergic neurons at F1 = 0.763.

## Extended DM Graph Reconstruction Algorithm

Although for interpretive purposes, we want to compute the unstable 1-manifolds (mountain ridges) of the Jaccard index, it is computationally easier to compute stable 1-manifolds (valley ridges). Stable 1-manifolds of the inverted Jaccard index are what is explicitly computed in the *Extended DM Graph Reconstruction Algorithm* but is intuitively analogous toto the unstable 1-manifolds of the original function.

***Algorithm:* Extended DM Graph Reconstruction Algorithm** (*X*, *k*, *δ*)

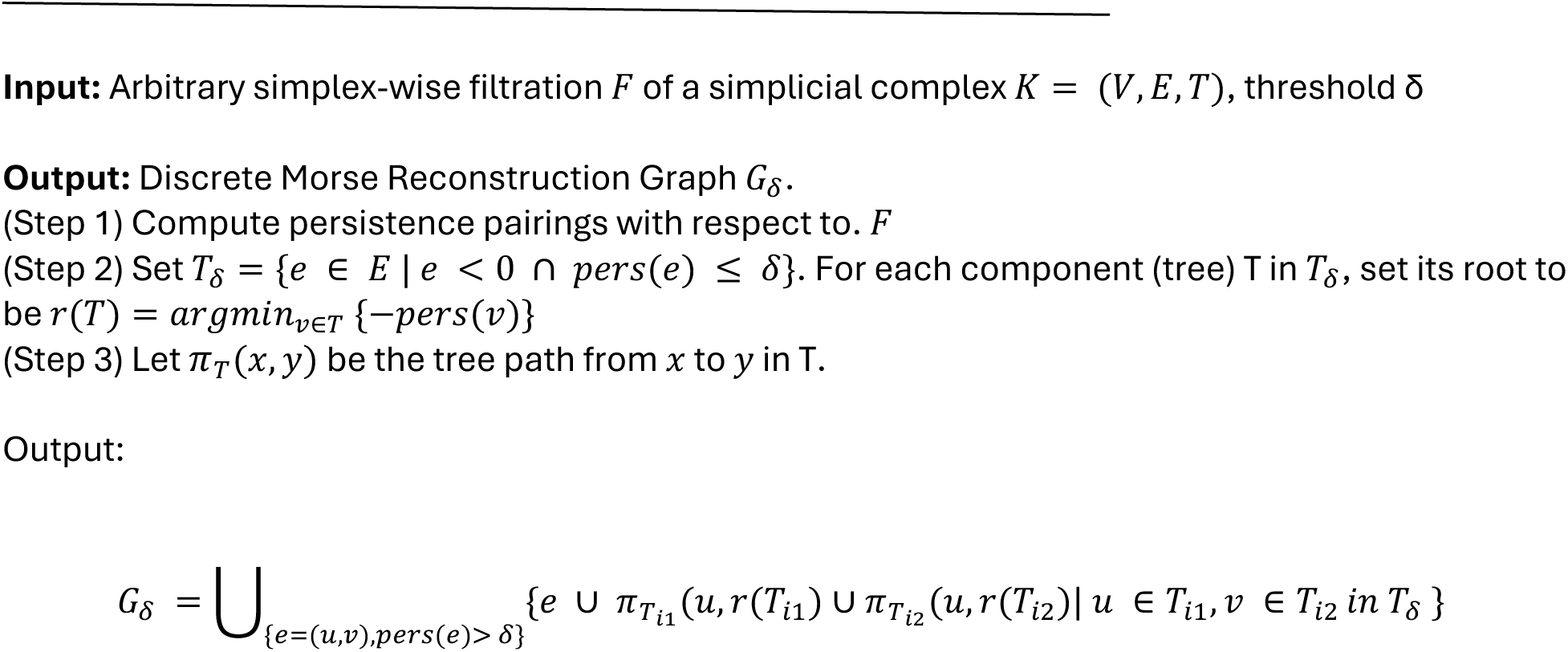

